# Kap-centric Nsp1-mediated nuclear transport at full amino-acid resolution

**DOI:** 10.1101/2024.06.23.600256

**Authors:** Maurice Dekker, Hendrik W. de Vries, Koen A. Wortelboer, Harm Jan Beekhuis, Maarten G.J.E. van Oosterhout, Erik Van der Giessen, Patrick R. Onck

## Abstract

Recent studies of nuclear pore complexes (NPCs) have provided detailed descriptions of the core scaffold structures, yet fall short in resolving the dynamic FG-meshwork with similar precision. Here, we present a novel modeling framework that enables the simulation of nuclear transport at full amino-acid resolution. We describe the distributions of the different FG-Nups in the central transporter and highlight the dynamic nature of the FG-meshwork, with FG–FG interaction lifetimes on the order of nanoseconds. Our findings reveal that Nsp1, the most abundant FG-Nup in the NPC, creates a central meshwork due to its unique bimodal structure, that is essential for controlling both passive and active transport. By adding nuclear transport receptors (NTRs)—specifically Kap95—to the pore, we demonstrate that NTRs play a key role in increasing the energy barrier for translocation of inert particles. The NTRs are subject to a dynamic interplay between binding to FG motifs and the temporal fluctuations of the FG-meshwork, leading to transient voids through which they move. Overall, our simulations identify a dense GLFG-ring coated by lower-mobility Kaps and a central dynamic FG–FG meshwork to create a reduced-dimensional transport surface of optimal binding avidity that drives Kap translocation.

## Introduction

Nuclear pore complexes (NPCs) regulate the transport of molecules between the nucleus and cytoplasm, ensuring the precise localization of proteins, RNA, and other essential molecules within the cell [1]. These large pores play a pivotal role in maintaining cellular homeostasis, gene expression, and the proper functioning of eukaryotic cells [2]. The permeability barrier of NPCs is highly selective, with most large macromolecules able to efficiently traverse the pore only when bound to nuclear transport receptors (NTRs). NTRs possess multiple binding sites that interact with phenylalanine–glycine (FG) motifs, which are abundant within the intrinsically disordered FG-meshwork that fills the central channel. NTRs are considered an integral component of the NPC and can dynamically modulate the pore’s microenvironment to enable fast and selective transport [3–6].

NPCs have been extensively studied using cryo-electron tomography and sub-volume averaging across various species, including humans [7–10], yeast [11–16] and algae [17]. These studies provide detailed descriptions of the core scaffold structures, yet fall short in resolving the highly dynamic FG-meshwork due to the dynamic nature of the FG-Nups and transport factors in the central transporter (CT). As a result, the functional organization underlying NPC barrier function and selective nuclear transport remains largely elusive. Super-resolution imaging techniques are able to track NTRs interacting with and transiting through the NPC [18–20], and recent advances using MINFLUX have now enabled high-precision tracking of single import and export events through individual NPCs at high spatiotemporal resolution [21]. However, these experimental approaches still lack direct information on the local FG-Nup structure and molecular interactions governing transport. To unveil the mechanisms underlying NTR translocation, molecular dynamics (MD) models have been developed that can act as a “computational microscope”, scrutinizing the molecular interactions at the scale of individual FG-Nups and NTRs, and achieving spatial and temporal resolutions that are inaccessible to experiments.

Atomistic MD simulations are a potentially powerful tool to study the key features of nuclear transport on a molecular level, yet the sheer size and complexity of the NPC and its cargo-complexes limits the applicability of such detailed models to short time and length scales [10, 22]. In order to mitigate these limitations, simplified coarse-grained molecular dynamics (CGMD) models have emerged [23–29], albeit at the cost of smearing out molecular detail and losing the amino-acid footprint characteristic of the highly heterogeneous collection of FG-Nups and NTRs. Building on these advances, we have developed a CGMD model with amino-acid resolution that captures the precise amino-acid sequences of individual FG-Nups, while also encompassing the collective behavior of all FG-Nups in the NPC at experimental timescales [30, 31]. This model has been used by us [32–39] and others [40–42] to study the collective FG-Nup density distribution inside the pore, albeit with rather simplified cargo or Kap models. A comprehensive simulation framework for nucleocytoplasmic transport at amino-acid resolution has been lacking until now, which has hampered a fundamental understanding of the molecular mechanisms underlying selective nuclear transport.

In this work, we present a complete residue-scale model of the yeast NPC, encompassing all 216 FG-Nups with lengths ranging from 250 to 800 residues, diverse amino-acid sequences, and proper protein stoichiometry. Our model incorporates an amino-acid resolution structural scaffold with FG-Nup anchor site locations obtained from the structural model, and, importantly, a large ensemble of Kap95 nuclear transport receptors represented at the same resolution. Large-scale, highly-parallel computations with the model highlight the essential role of Nsp1 in establishing a dynamic permeability barrier through the formation of a dynamic, low-density percolation at the center of the pore, facilitated by highly dynamic FG–FG cross-links with nanosecond interaction lifetimes. This barrier not only regulates the passage of inert cargo in a size-selective manner but also facilitates efficient transport through its distinctive FG-rich extended domains that span across the center of the pore. Our findings further underscore the critical role of transport factors in maintaining the NPC’s barrier function. Increasing the concentration of Kap95 reinforces the selective properties of the permeability barrier, consistent with “Kap-centric” models [3–6]. This enhanced selectivity arises from two effects. First, the accumulation of Kaps within the CT increases the effective energy barrier for passive translocation of inert molecules. Second, our simulations show that NTR-mediated transport is a discontinuous process, in which temporarily slow Kaps decorate the sticky surface of the dense GLFG-ring, thereby facilitating the rapid translocation of other Kaps along the interface between the GLFG-ring and the Nsp1 percolation. Together, these findings reveal a dynamic transport mechanism in which Kaps traverse the NPC through transient voids, characterized by the interplay between FG motif binding and steric hindrance.

Our residue-scale computational approach lays the foundation for multiple future directions in follow-up work, including dilation and constriction [7, 13], structural and compositional variations [7, 12, 14, 16], simultaneous import and export [20, 21, 29], and cross-species comparison [17, 21, 43, 44], opening the exciting opportunity to fully resolve the essential physical principles of selective nuclear transport at unprecedented molecular detail.

## Results

### The yeast NPC at amino-acid resolution

We developed a computational model of the yeast NPC at amino-acid resolution, based on the integrative structure of Kim *et al*. [11]. The model employs our previously developed one-beadper-amino-acid (1BPA) force field for disordered FG-Nups [30, 31, 34–36, 38, 39], which includes residue-specific electrostatic, hydrophobic and cation–π interactions for all twenty amino acids, as well as backbone potentials that capture the local stiffness of the chain by distinguishing between glycine, proline and other residues. Simulations were performed using an implicit solvent representation, allowing efficient sampling of large systems. Our model incorporates the full scaffold structure from Ref. [11] at near-atomic resolution (Fig. 1a), by placing amino acid beads at each C_α_ position (Fig. 1b). A torus-shaped occlusion is included to represent the nuclear membrane (Methods). Undetermined regions in the scaffold Nups and the initial configurations of the FG-Nup meshwork are generated using a biased self-avoiding random walk (Fig. 1c, Methods). The anchoring locations for each of the disordered FG domains were selected based on the data in Ref. [11] (Fig. 1d,e). As the reference structure lacks sufficient resolution for the nuclear basket proteins Mlp1 and Mlp2 and the corresponding anchoring site of Nup2, these components are not included in the current model. The complete model of the yeast NPC at full amino-acid resolution is illustrated in Fig. 1f.

**Fig. 1.**
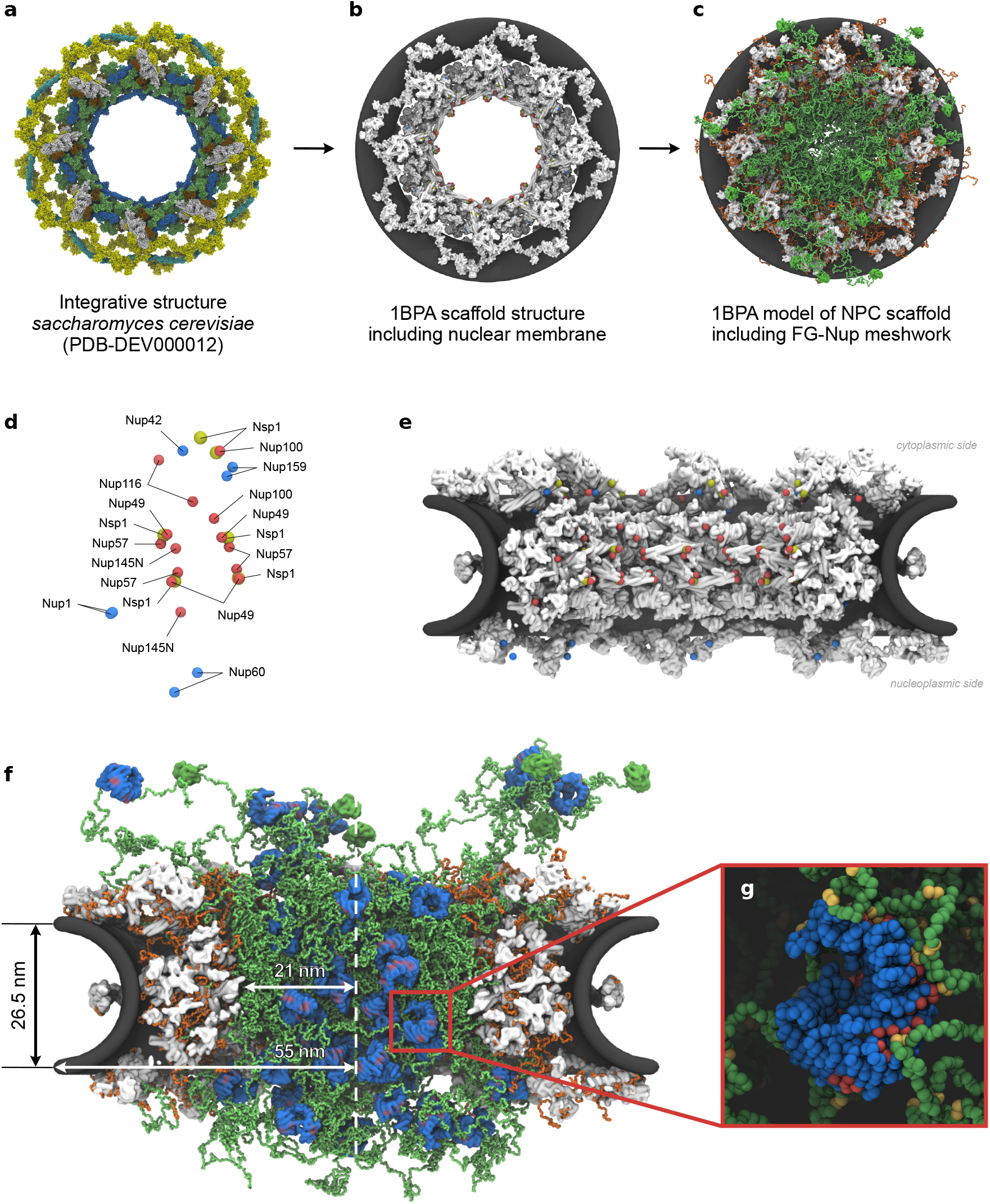
Residue-scale model derived from an integrative structure of the yeast NPC. (**a**) All-atom reference structure of the yeast NPC [11]. The scaffold structure, membrane placement and disordered FG-Nup anchoring coordinates in our residue-scale model are all derived from this structure. The coloring scheme follows a grouping as defined in the original study [11]. (**b**) Residue-scale scaffold structure (white), including a torus-shaped occlusion (dark gray) that mimics the nuclear membrane. GLFG-Nup anchoring sites are indicated in red, FG-Nup anchoring sites are indicated in blue (FG-Nups: Nup159, Nup42, Nup1, Nup60) and Nsp1 anchoring sites are yellow. Note that not all anchoring sites are visible in this orientation due to the geometry of the scaffold. (**c**) A self-avoiding pseudo-random walk is used to generate missing loops in the scaffold structure (orange) and configurations of disordered FG-Nup segments (green), thus completing the residue-scale model of the yeast NPC. (**d**) Anchoring sites of individual GLFG-Nups (red), FG-Nups (blue) and Nsp1 (yellow) for a single spoke, including stoichiometry. (**e**) FG/GLFG-Nup anchoring sites (side-view), highlighting four out of eight symmetric NPC spokes. (**f**) Side-view and geometric details of the single-residue computational model of the yeast NPC including NTRs (Kap95, blue), following the coloring scheme of (c). The folded domains in the FG-Nup meshwork (green surface structures) are the β-propellers at the N-termini of the Nup159 FG domains. See Suppl. Movie 1 for a short trajectory of the Kap and FG-Nup dynamics. (**g**) Zoom in on the amino-acid details of a single Kap95 protein (blue) with binding sites (red) in the FG-meshwork (green) featuring FG motifs (yellow). Not shown: the blue beads include cationic, anionic and aromatic residues, while the green beads include all 20 residues.

### Structural organization of FG-Nups in the CT

We began by simulating the NPC in the absence of translocating particles. After an extensive equilibration of the FG-meshwork (Methods), we conducted an equilibrium simulation of 2.5 ms that is used for analysis. We first analyzed the relative localization of the different FG-Nup species in the CT. In the equilibrium state, all GLFG-Nups (Nup116, Nup100, Nup145N, Nup49, and Nup57) coassemble into a high-density, ring-like structure at the center of the NPC (green in Fig. 2). This “GLFG-ring” forms a high-density barrier that reduces the effective diameter for passive transport (Fig. 2f and Suppl. Fig. S4).

**Fig. 2.**
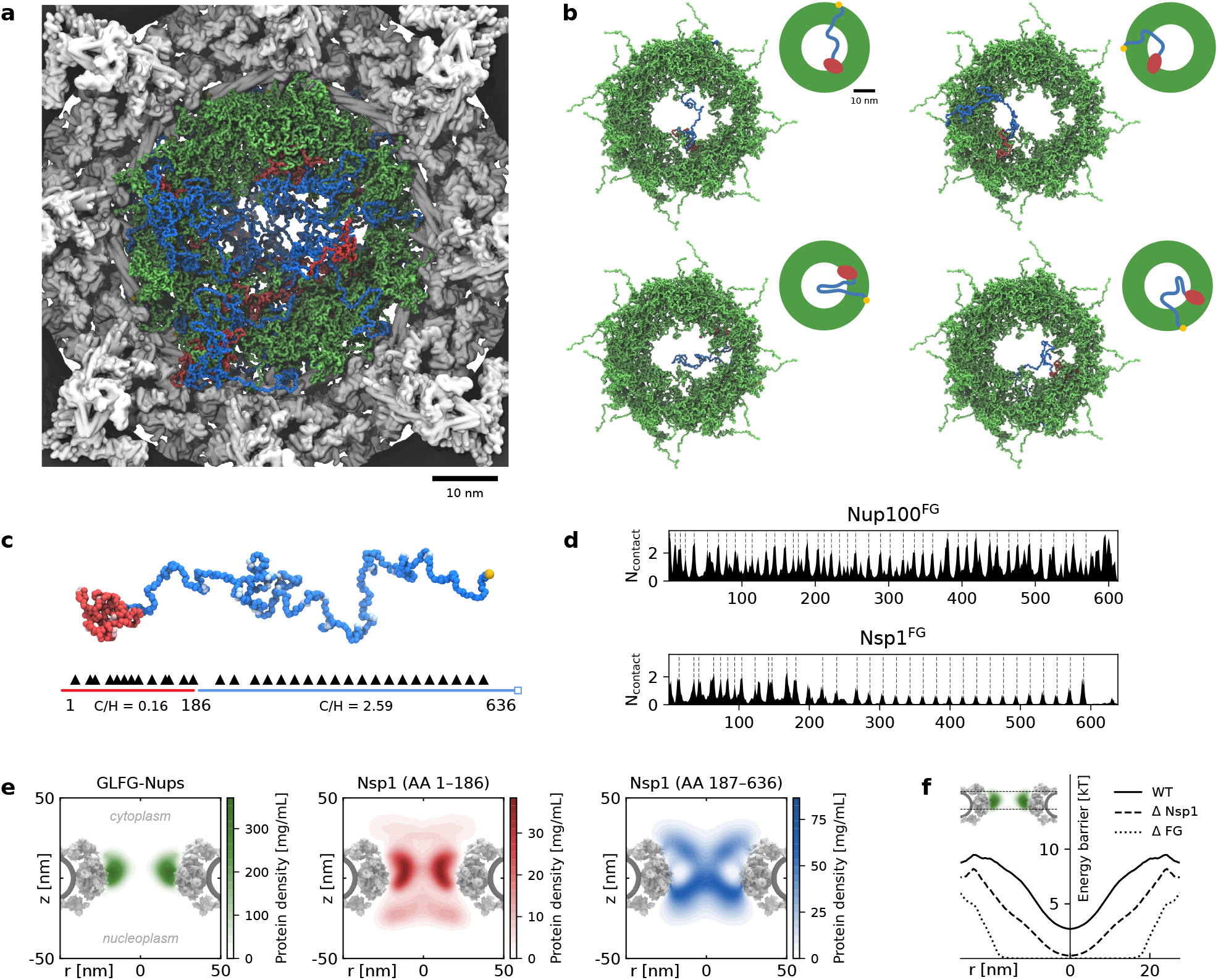
The dynamic permeability barrier of the NPC. (**a**) Top view of the pore’s central channel. The GLFG-Nups (green) form a ring-like assembly near the pore’s scaffold. The center of this GLFG-condensate is filled by a low density percolation, formed mainly by the extended domains of Nsp1 (blue). Nsp1 molecules are anchored to the scaffold at their C-terminal (yellow), while the cohesive N-terminal domains (red) interact with the GLFG-Nups. Cytoplasmic and nucleoplasmic Nups are not shown. (**b**) Examples of individual Nsp1-proteins in the center of the pore. In all cases, the extended domain protrudes from the GLFG-condensate towards the center of the pore, while the collapsed domain sticks to this GLFG-condensate. (**c**) Bimodal structure of Nsp1 highlighted by a cohesive collapsed domain (red) and a noncohesive extended domain (blue). FG motifs in the respective domains are highlighted by light red and blue beads; the anchor to the NPC scaffold is highlighted in yellow. The relative location of FG motifs along the Nsp1-sequence is shown by the black markers below the protein structure. (**d**) Time-averaged number of intermolecular contacts of the FG domains of Nup100 and Nsp1 with all other Nups in the NPC (FG domains only). The dashed lines indicate the location of FG motifs. Intermolecular contact maps for all other FG-Nup species in the pore are shown in Suppl. Fig. S3. (**e**) Radially-averaged density distributions of the GLFG-Nups and both domains of Nsp1, demonstrating that the cohesive N-terminal domain of Nsp1 primarily localizes around the GLFG-ring. (**f**) The energy barrier for passive translocation of inert particles of radius *r*_p_ = 3.4 nm at the center of the pore for the full NPC (WT), when Nsp1 is removed (Δ Nsp1) and when all FG domains are removed (Δ FG). The inset of the NPC scaffold with GLFG-Nups shows the region that is used for analysis. Note that the analysis considers the same simulation trajectory in all three cases; the Nup deletions are done during post-processing.

With 48 copies, Nsp1 is the most abundant FG-Nup in the NPC [11]. In the CT, Nsp1 molecules are the only FG-Nups spanning the opening left by the GLFG-ring. The C-terminus of Nsp1 is tethered to the scaffold wall, while the cohesive, collapsed N-terminal domain dynamically coassembles with the Nups in the GLFG-ring (red in Fig. 2). Consequently, the extended domains of Nsp1, which uniquely feature a high C/H ratio and numerous FG motifs [38, 45] (Suppl. Fig. S1), span across the central opening of the GLFG-ring (blue in Fig. 2). The eight-fold rotational symmetry of the NPC scaffold results in Nsp1 proteins spanning the CT from multiple directions, creating a low-density yet stable percolated network that efficiently blocks large inert cargoes from translocating across the NPC. Indeed, removal of Nsp1 from the CT significantly lowers the energy barrier for passive translocation (Fig. 2f and Suppl. Fig. S4).

The FG-Nup assembly in the CT is primarily stabilized by hydrophobic FG–FG interactions (Fig. 2d), a characteristic also observed in FG-Nup condensates [38]. These interactions are highly dynamic, with lifetimes on the order of nanoseconds (Suppl. Fig. S5), similar to the transient cross-links found in FG-Nup condensates. Importantly, the measured lifetimes are comparable across all FG-Nup types (Suppl. Fig. S6), indicating that the dynamics of FG–FG interactions are largely independent of Nup identity. Interestingly, the composition and organization of the GLFG-ring observed here closely resemble that of the GLFG-condensate studied previously [38].

### Kap95 proteins coat the GLFG-ring

It has been shown that NTRs can significantly enhance the selectivity of the permeability barrier [6]. However, on a molecular level, the conformational changes of the FG-Nups in response to high concentrations of NTRs remain largely elusive. To elucidate how NTR localization within the NPC affects the selective permeability barrier, we performed a computational titration experiment by systematically increasing the concentration of Kap95 proteins in the CT.

To minimize the re-equilibration time upon introducing a substantial number of translocating particles into the FG-meshwork, we developed a custom pipeline to insert Kap95 proteins into the FG-meshwork by “growing” them at randomly assigned coordinates inside the CT (see Suppl. Methods for more details). We generated five NPC–Kap95 systems with strongly increasing numbers of Kap95 proteins, ranging from a system without Kaps to a system where the FG-meshwork is fully saturated by the Kaps (Fig. 3). This approach allows us to compare the equilibrium dynamics in the CT across a wide range of Kap95 concentrations. From the equilibrium trajectories, we calculated the time-averaged density distributions of the FG-meshwork and the Kap95 proteins (Fig. 3d–f). Notably, we find that the NPC can accommodate a remarkable number of Kap95 proteins; at the highest concentration the NPC can no longer fully accommodate all Kap95 proteins, with up to 120 Kaps in contact with the FG-meshwork. Furthermore, at all Kap95 concentrations, the Kaps primarily localize near the high-density GLFG-ring, effectively “coating” the GLFG-assembly (Fig. 3f). These interaction regions are consistent with super-resolution microscopy studies in human NPCs [18, 19] and have been associated with the translocation pathway of importins. While Kap95 proteins tend to occupy regions with a high concentration of FG motifs, their localization is constrained by the high density of the GLFG-ring. As a result, rather than entering the GLFG-assembly, the Kap95 proteins reside around the high-density GLFG-ring where there is a high local FG motif density but low overall protein density. Increasing the local Kap95 concentration results in a layer of Kaps around the cohesive GLFG-ring, occupying most of the sticky FG motifs of the GLFG-Nups. As a result, other NTRs experience weaker FG-interactions and become more dynamic, facilitating easier translocation across the NPC [46], mainly by interacting with the central percolation formed by the extended Nsp1 domains. At high Kap95 concentrations, the Kap density distributions (Fig. 3f) resemble the hourglass-shaped central “plug” that is often reported by cryo-EM experiments [11, 14, 16].

**Fig. 3.**
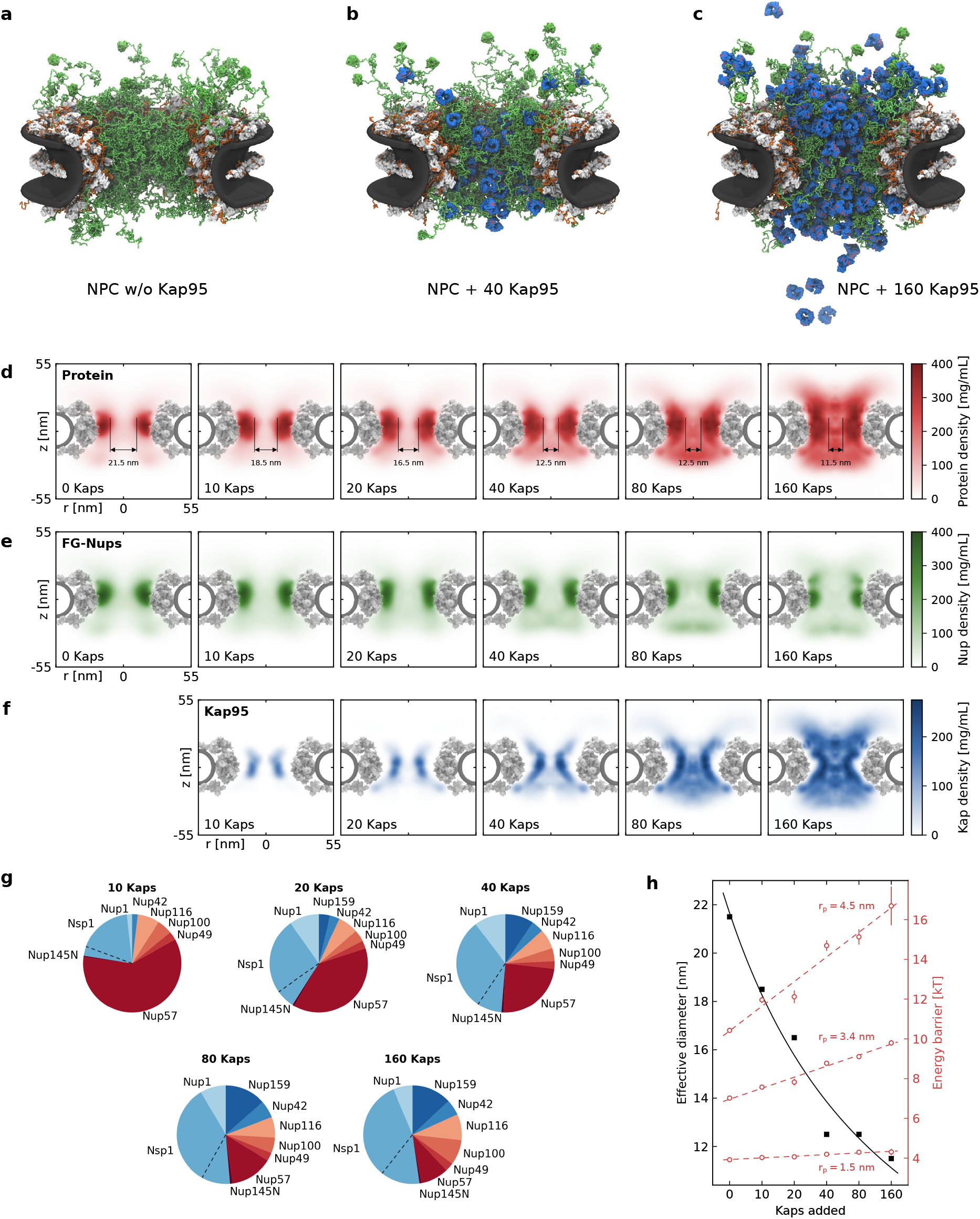
Effect of Kap concentration on the FG-Nup meshwork in the NPC. (**a–c**) Simulation snapshots of the yeast NPC (a) in the absence of Kap95, (b) with 40 Kap95 proteins, and (c) with 160 Kap95 proteins. (**d**) Radially-averaged distribution of the total protein density (Nup and Kap95) in the NPC at varying concentrations of Kap95. With increasing concentration Kaps start coating the GLFG-ring, thus effectively decreasing the diameter of the low density region in the center. (**e**) Radially-averaged FG-Nup density distribution in the NPC at varying concentrations of Kap95. (**f**) Radially-averaged Kap95 density distribution in the NPC at varying concentrations of Kap95. (**g**) Relative fractions of FG-Nup–Kap interactions by FG-Nup type. At low Kap concentrations, the Kaps primarily interact with the GLFG-Nups (red). At higher Kap concentrations Kaps interact mainly with the other Nups (blue). The dashed lines mark the relative components of the collapsed (bottom) and extended (top) domain of Nsp1, respectively. (**h**) Kap95 crowding reduces the effective diameter of the (passive) transport channel resulting in a high selectivity for large inert cargoes. The effective diameter of the transport channel as a function of the number of Kaps added to the NPC (black squares) and the energy barrier for (passive) translocation for particles with a radius of 1.5 nm (e.g. ubiquitin), 3.0 nm (e.g. BSA) and 4.5 nm (red circles).

### Kap95 crowding enhances NPC selectivity in a size-selective manner

Next, we asked how the presence of Kaps influences the NPC permeability barrier. Initially, in the absence of Kap95 proteins, the average FG-occupancy—defined as the percentage of FG motifs cross-linking with another FG motif—is approximately 40 %, similar to FG-Nup condensates [38]. Surprisingly, the addition of Kap95 proteins does not significantly alter the number or distribution of FG–FG cross-linking interactions present in the NPC, consistent with Ref. [37]. Instead, we find a constant number of FG–FG cross-links, independent of the Kap95 concentration, occurring largely in the same regions (Suppl. Figs. S9a and S10). This constancy is explained by the relatively low FG–Kap occupancy (*<*30 %) and the presence of a dynamic, unbound FG motif pool (20 %; Suppl. Fig. S11). A breakdown by Nup-type shows that GLFG-rich Nups dominate FG–FG interactions, although FxFG-Nups, including Nsp1, also contribute substantially (Suppl. Fig. S12). A closer look at the density distribution of the FG-Nups confirms that the FG-meshwork is largely unaffected by Kap95 binding (Fig. 3e), suggesting that Kaps do not compete with FG motifs, but rather bind to the large pool of still available FG motifs.

To determine the effect of Kap95 crowding on passive transport, we calculated the energy barrier for translocation of inert cargoes of varying sizes as a function of Kap95 concentration (Fig. 3h). We found that for small cargoes (*r*_p_ = 1.5 nm, e.g. ubiquitin) there is only a small increase in the energy barrier for translocation. For larger cargoes (*r*_p_ = 3.4 nm, e.g. BSA) the energy barrier for translocation increases with Kap95 concentration.

For the simulations with 80 and 160 Kaps (and to a lesser degree also the system with 40 Kaps) we noticed an accumulation of Kap95 proteins at the nuclear face of the NPC. Even at these high Kap concentrations, only a few Kaps have dissociated from the FG-meshwork into the nucleoplasm. Instead, the Kaps at the nuclear face of the NPC form a cross-linked mesh together with Nup1. This is consistent with experimental observations of human NPCs where an accumulation of Nup153 (the homologue of Nup1) at the nuclear side has been observed when RanGTP is absent [47, 48].

### Kap translocation is driven by FG-interactions and dynamics of the meshwork

To mimic the effect of the Ran-gradient, we have repeated the simulation with 80 Kaps, this time recycling the Kaps that reach the nuclear face of the pore (see Methods for details on the procedure). This recycling procedure induces a moderate flux of Kap95 proteins through the pore, allowing us to study the translocation pathway of Kaps in more detail. Our simulations span a timescale comparable to the millisecond duration of Kap transport events [49, 50], providing valuable insights into the translocation dynamics of Kaps within the CT.

In contrast to inert molecules, Kap95 proteins have a high affinity towards the FG-meshwork through the numerous available FG motifs within the permeability barrier. In the previous section, we observed that Kaps tend to populate the regions rich in FG motifs. Due to the frequent, nanosecond-long contacts, many Kaps are dynamically bound to the surface of the central GLFG-assembly, potentially impeding their diffusive speed. Despite this, the Kaps are not stuck; several Kap95 proteins translocate across the permeability barrier, albeit with largely varying translocation times. An example of such a translocation event is shown in Fig. 4 and Suppl. Movie S2. It is remarkable to observe that the Kaps do not move at a constant speed. Instead, they are predominantly confined within their local environment and traverse the pore through many small “jumps” to nearby locations. This confinement is primarily caused by the steric hindrance of the local FG-meshwork and the multivalent FG–binding interactions. Therefore, the translocation of Kaps requires two conditions: a temporary reduction of FG-binding interactions and the presence of openings (“voids”) in the meshwork through which the Kap can diffuse. The translocation of Kaps across the CT can be described as a cascade of small random steps without a pronounced overall directionality. The overall directionality is predominantly described by the Ran gradient, off-loading the Kaps when they reach the nuclear face of the pore [51].

**Fig. 4.**
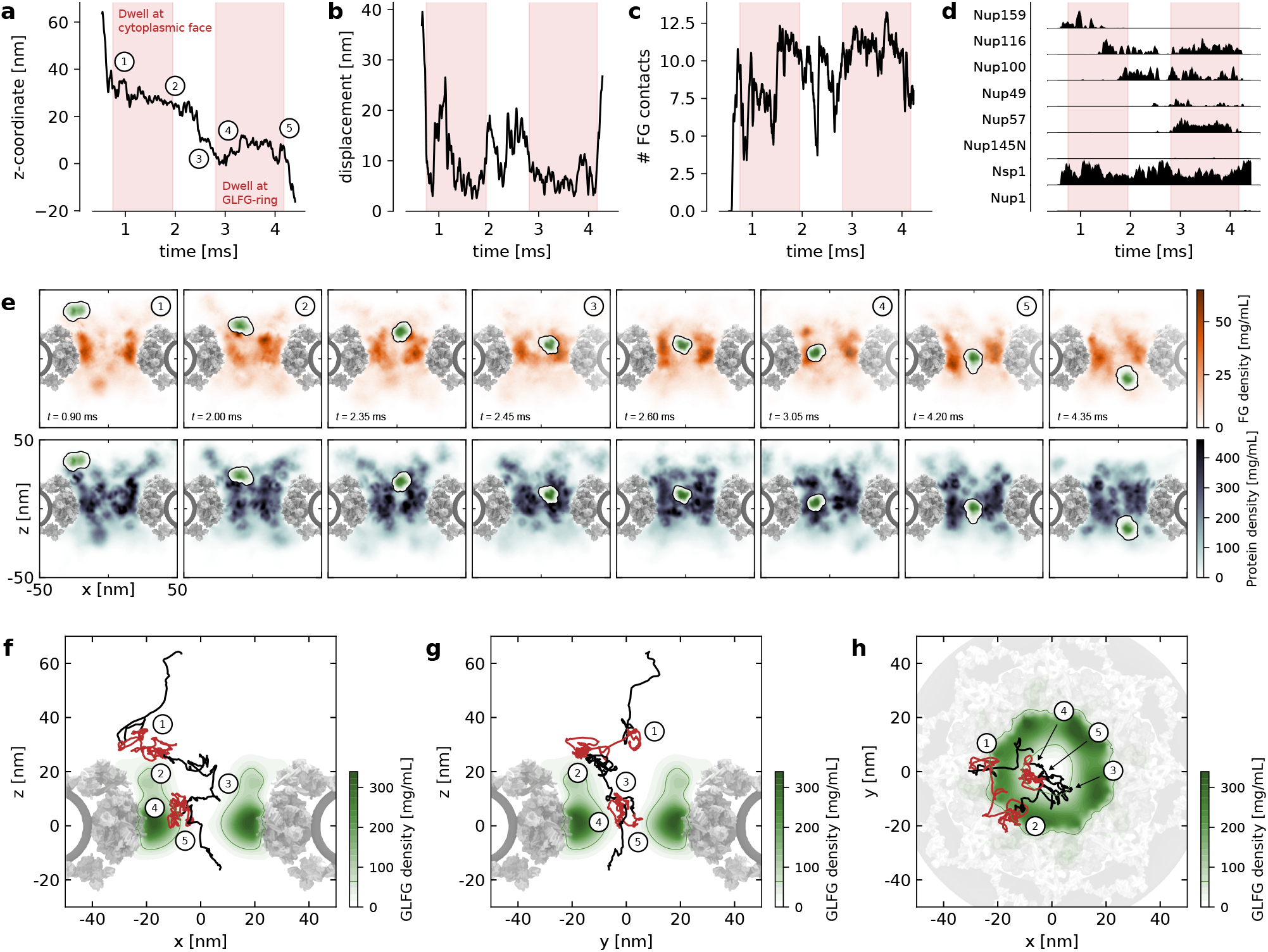
Kap dynamics is described by FG motif density and available space (steric hindrance). (**a**) Position of a selected Kap along the pore axis as a function of time. The Kap does not translocate in one go, rather it dwells at several locations in the pore (red time intervals). (**b**) Displacement of the selected Kap as function of time, where the displacement is measured over a time window of 250 µs. (**c**) Number of FG-interactions that the selected Kap experiences as a function of time. (**d**) Contacts per FG-Nup type. (**e**) Selected simulation snapshots of the selected Kap overlaid on top of the FG motif distribution (top row) and the total protein distribution of both FG-Nups and Kaps (bottom row). Each snapshot represents an averaged density projection over a 20 nm thick slice of the simulation volume, centered on the Kap and taken in the *x* –*z* plane (in contrast to the *r* –*z* projections used elsewhere). The complete translocation event is shown in Suppl. Movie S2. (**f–h**) Trajectory of the selected Kap shown on top of the average density distribution of the GLFG-Nups. The red segments of the trajectory correspond to the time intervals highlighted in panels (**a–d**). The trajectory is projected onto the (**f**) *x* –*z*, (**g**) *y* –*z*, and (**h**) *x* –*y* planes.

### Kaps traverse the pore in multiple discrete steps

A detailed description of a single Kap translocation event (Fig. 4) illustrates this process. Initial contact with the NPC occurs through interactions with the peripheral Nup159 and Nsp1 (Fig. 4d). At this location, the FG-meshwork has a relatively low density, allowing for considerable mobility. Due to the crowding of other Kaps at the cytoplasmic face of the pore (Fig. 3f), the Kap is partially screened from the sticky FG-interactions of the GLFG-ring, allowing dynamic movement during the first half of this stage. As the Kap progresses, it encounters the cytoplasmically anchored Nup116 (and later also Nup100) (Fig. 4d), leading to an increase in FG contacts and a subsequent reduction in mobility. The Kap becomes locally confined and dwells for some time at the cytoplasmic face of the pore (location 2).

We draw particular attention to the density fluctuations of the FG meshwork, as shown in Fig. 4e, which play a crucial role in Kap translocation. As the Kap moves from location 2 to 3, it is handed over to the opposite side of the pore, where the FG motif density is high as well and the total density is temporarily low (*t* = 2.0 ms, bottom row of Fig. 4e). Subsequently, an opening (void) appears in the center of the pore (*t* = 2.35 ms, bottom row of Fig. 4e), allowing the Kap to slide downwards over the GLFG surface (2.35 *< t <* 2.45 ms, top row of Fig. 4e). Then, from location 3 to 4, the Kap moves to the left side of the GLFG-ring and continues downward, arriving at location 4. Additional views of the trajectory from multiple angles further confirm that the Kap traverses the central pore volume (Fig. 4f–h).

In the final phase, the Kap gets temporarily stuck on the GLFG-ring (location 4) due to numerous additional interactions with Nup57 (and Nup49), while interactions with Nsp1 are slightly reduced (Fig. 4d). This reduced mobility lasts until the interactions with Nup57 are disrupted at location 5, and the Kap detaches from the GLFG-ring (*t* = 4.20 ms, top row of Fig. 4e). Finally, nonetheless guided by Nsp1, the Kap diffuses to the nuclear face and is “removed” from the CT by RanGTP. Notably, although the Kap is continuously in contact with Nsp1 chains during translocation, the translocation pauses do not correspond to peaks in Nsp1 interactions. Instead, these dwell phases align with increased engagement with GLFG-Nups (Fig. 4d). This detailed observation of a single Kap translocation event highlights the intricate and dynamic nature of nuclear transport, underscoring the crucial role of FG motifs and the dynamic Kap-containing FG-meshwork in facilitating efficient translocation through the NPC.

A comprehensive set of additional translocation events is provided in Suppl. Figs. S17–S23, illustrating that the dynamic features observed in the main text are representative of multiple Kap95 translocations.

### “Dirty velcro” mechanism is essential for efficient translocation

To gain a more comprehensive view of the translocation process, we analyzed all eight complete Kap95 translocation events observed in our recycling simulation (Fig. 4 and Suppl. Figs. S17– S23). When projected in *r* –*z* space on top of the GLFG-ring density distribution, these trajectories reveal substantial variation in both pathway and translocation speed, yet a consistent trend emerges: Kaps predominantly migrate along the surface of the GLFG-ring rather than through the central axis of the pore. This observation is in line with recent single-molecule tracking experiments in human NPCs [21], which similarly reported that transport trajectories avoid the central channel and are instead confined to more peripheral pathways within the pore.

To further illustrate how interactions with the GLFG-ring influence the mobility of translocating Kaps, we compared the spatial distribution of their slow and fast phases (Fig. 5b). These density distributions show that the highest Kap mobility occurs at the cytoplasmic face of the pore, where the FG-meshwork is relatively sparse and Kaps primarily interact with Nup159 and Nsp1 (Fig. 4d). The lower-mobility Kaps are predominantly localized along the surface of the GLFG-ring, where their dynamics are reduced by the many FG–binding interactions. Notably, these slower Kaps effectively screen other Kaps from the sticky FG-interactions of the GLFG-ring, thereby facilitating more rapid transport toward the pore center. This suggests that while the FG-meshwork promotes initial Kap engagement and mobility, the denser GLFG regions act as dynamic checkpoints, temporarily slowing down some Kaps to enable faster passage of others through reduced FG-binding [50, 51].

**Fig. 5.**
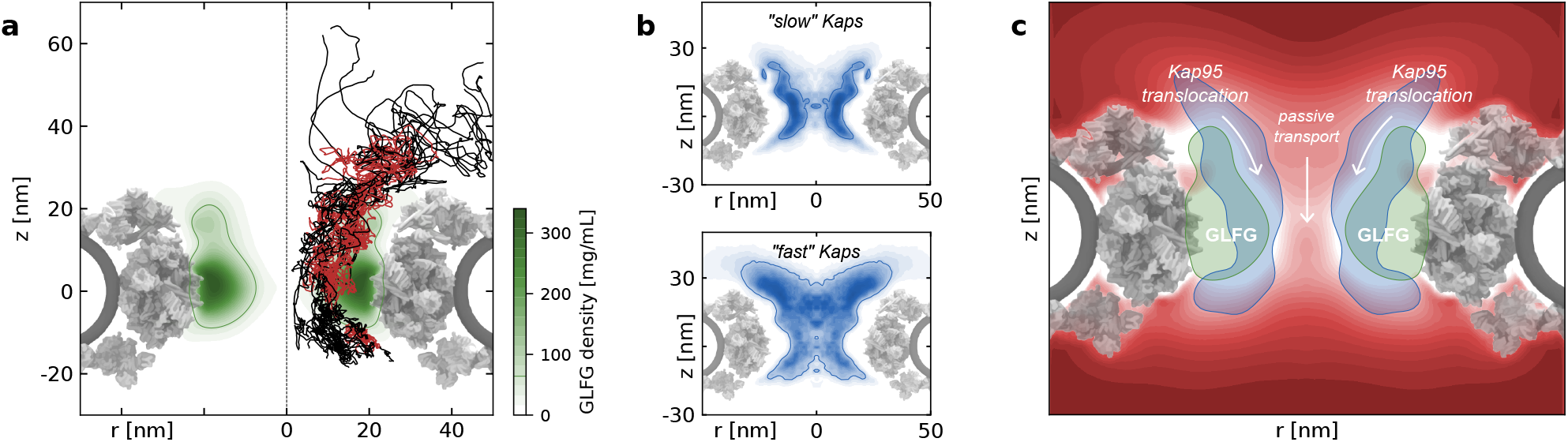
Transport mechanism in the yeast NPC. (**a**) All eight complete Kap95 translocation trajectories from the recycling simulation (Fig. 4 and Suppl. Figs. S17–S23), projected in *r* –*z* space and overlaid on the average density distribution of the GLFG-Nups (green). The trajectories are color-coded as in Fig. 4, with black indicating dynamic movement and red indicating dwell phases. (**b**) Spatial distribution of the slow phase and fast phase of Kap displacements, highlighting regions of low and high mobility, respectively. (**c**) Schematic representation of the proposed translocation mechanism. GLFG-Nups form a dense, ring-like structure at the center of the pore (green). Kaps translocate through the NPC via a reduction-of-dimensionality mechanism, moving predominantly along the GLFG-ring surface. Kaps that are temporarily bound to this surface form a coating that shields other Kaps from strong FG interactions, enabling faster movement along the interface between the GLFG-ring and the dynamic Nsp1 network. In contrast, passive translocation of small inert molecules (red) occurs primarily through the central Nsp1 percolation pathway (Fig. 2f and Suppl. Fig. S13). Note that the CT is highly dynamic; average density distributions do not show the complete picture.

## Discussion

### Dynamic architecture of the CT highlights the important role of Nsp1

We have developed a residue-scale MD framework for nuclear transport in the yeast NPC to study the conformational changes of the permeability barrier during NTR translocation. We used this framework to study the yeast NPC under drastically varying Kap95 concentrations, starting from an NPC devoid of NTRs. Our simulations revealed a large variation in protein density in the CT. Specifically, the cohesive GLFG-Nups form a ring-like assembly at the scaffold surface, that is sustained by numerous, highly dynamic FG–FG cross-links. The opening left by this GLFG-ring is filled with a low-density, percolated network of Nsp1 proteins. Due to their unique bimodal nature—containing a 200-residue cohesive domain together with a 400-residue noncohesive, extended domain, both encompassing many FG motifs—Nsp1 proteins are able to span across the central pore opening. Their C-terminal is anchored to the NPC scaffold, while the cohesive N-terminal domain dynamically co-assembles with the GLFG-Nups that line the inner scaffold surface. The many FG motifs in the extended domains of Nsp1 establish a cross-linked yet dynamic percolation at the center of the pore, acting as a size-selective filter for inert molecules [42].

The GLFG-ring at the center of the CT has multiple functions. First, it acts as a seal between the pore scaffold and the central Nsp1 network, preventing leakage of macromolecules along the scaffold surface. Second, it provides a dynamic anchoring site for the cohesive Nsp1 domains, enabling their extended domains to span the central channel. This anchoring is essential for maintaining the Nsp1 percolation at the pore center, as its absence leads to the expulsion of Nsp1 from the pore [37, 52].

In addition, Nsp1 chaperones Kaps throughout their entire translocation pathway, from initial capture at the cytoplasmic side to final release at the nuclear side (Fig. 4d). The FG motifs in the extended domain of Nsp1 are crucial for this process, as removing only the FG–Kap95 interactions in these domains effectively eliminates all transport (Suppl. Fig. S15). It is noteworthy that the extended domain of Nsp1—being the only FG domain in yeast that combines low cohesivity with regular FG motif spacing—plays such a pivotal role in nuclear transport.

In this context, we note that Hülsmann *et al*. [53] ascribed a central role to Nup98 in human NPCs and concluded that cohesive domains alone are sufficient to form a permeability barrier. While their study involved a reconstituted, heterogeneous system, they also found that any long, cohesive FG-Nup could restore barrier function, including those unrelated to Nup98. Our findings agree in that cohesive FG domains (e.g., in GLFG-Nups) are key for forming a physical barrier. However, in the yeast NPC, we find that the cohesive domains alone are insufficient for efficient transport: without FG-binding interactions from the noncohesive Nsp1 domains, translocation is strongly reduced (Suppl. Fig. S15). This indicates that, in yeast, it is the combination of the GLFG-ring and the dynamic, extended Nsp1 domains that creates a functional permeability barrier. Nsp1 is therefore not merely a space-filling component but a critical, active contributor to selective transport.

We propose that the dynamic nature of FG–FG cross-links in the GLFG-ring and Nsp1-network increases the robustness of the permeability barrier, allowing it to adapt to varying pore diameters [7, 13, 14] and accommodate large cargoes [28, 54–56] without compromising selectivity.

### Central transporter resembles an FG phase condensate

Experimental and computational studies have shown that the FG domains of several FG-Nups phase separate and form biomolecular condensates [38, 57–60] that are stabilized by a high degree of dynamic FG–FG cross-linking [38]. Our observations reveal an intriguing resemblance between these FG-Nup condensates and the high-density GLFG-ring in the CT. Both assemblies exhibit a highly dynamic percolated network with a similar degree of FG–FG cross-linking and similar cross-link dynamics. These properties remain largely unaffected by the partitioning of Kap95 in the CT, with only minor differences observed in the distribution of FG–FG cross-links between NPCs with and without Kaps.

Despite the extensive cross-linking, the FG-meshwork remains highly dynamic due to the characteristic “stickers-and-spacers” [61] architecture of FG-Nups. Cohesive FG motifs (stickers) drive the formation of a percolated network at the center of the pore, while FG-spacers (primarily consisting of polar and charged amino acids) mediate the contacts among these stickers [38, 60, 62]. This results in a dynamic FG-meshwork where the hydrophobic FG motif interactions are modulated by the hydrophilic character of the FG-spacers.

Although the ring-like GLFG-assembly cannot be strictly termed a phase condensate—since the FG-Nups cannot freely diffuse due to their tethering to the NPC scaffold [63, 64]—the similarity between the NPC’s GLFG-ring and FG-Nup condensates is striking. In our previous work [38], we studied the phase separation of an FG-Nup mixture resembling the stoichiometry of the NPC. Here, the central GLFG-Nups (Nup100, Nup116, Nup145N, Nup49 and Nup57) undergo co-condensation and form one large GLFG-Nup condensate, while the peripheral FG-Nups (Nup159, Nup1 and Nup60) exclusively remained in the dilute phase. Interestingly, a small fraction (10 %) of Nsp1 was found to co-condensate with the GLFG-Nups as well, primarily interacting through its cohesive N-terminal domain. This isolated “NPC condensate” has a remarkable resemblance with the organization of the FG-meshwork in the NPC, wherein the GLFG-Nups dynamically co-assemble into a ring-like structure and where the cohesive N-terminal domain of Nsp1 co-assembles with the GLFG-Nups. Although the FG-meshwork inside the NPC is tethered to the pore scaffold, both assemblies have comparable protein densities of approximately 400 mg*/*mL, contain the same co-condensation of five GLFG-Nups (Nup100, Nup116, Nup145N, Nup49 and Nup57), and are stabilized by a large number of dynamic FG–FG cross-links. Regardless of all these similarities, the key distinction between FG condensates and the CT is the tethered nature of Nsp1, allowing its extended domain to play a decisive role in nuclear transport. Remarkably, Nsp1’s efficiency is directly linked to its amino-acid sequence, which features an almost perfect FG-spacing of 15 residues, separated by spacers with a higher C/H ratio than any other yeast FG domain [45], making it uniquely suited for this function.

### Nsp1 as a phase state regulator in the NPC

Nsp1 has been identified as a phase state regulator for FG-Nup condensates in vitro [65]. In those studies, the cohesive N-terminal domains of Nsp1 co-condensate with GLFG-Nups, while the extended domains are projected from the condensate surface. Our simulations reveal a similar behavior: the cohesive domains of Nsp1 dynamically co-assemble with the GLFG-ring, while the noncohesive extended domains are projected towards the center of the pore. This suggests that Nsp1 may also play a phase regulatory role in the NPC, keeping the FG-Nups in a more dynamic state. By doing so, Nsp1 prevents the potential aggregation of FG-Nups into rigid beta-structures, thereby contributing to the robustness and stability of the permeability barrier.

### Kaps reinforce the selective permeability barrier

Our results support the Kap-centric models, which posit that Kaps are essential components of the selective permeability barrier [3–6]. At low Kap concentrations, the Kap95 proteins primarily localize around the GLFG-ring, effectively reducing the pore’s diameter and thereby limiting the leakage of unwanted molecules. At higher Kap concentrations, more Kap95 proteins not only localize around the GLFG-ring but also disperse throughout the entire FG-meshwork. Interestingly, the availability of FG motifs is not the limiting factor; instead, the available space at the center of the pore becomes depleted [37]. With increasing concentrations of Kap95, the permeability barrier becomes more size-selective [66]. The energy barrier for passive translocation of large inert particles (greater than 3.0 nm) is significantly increased, while the passive translocation of small inert particles (less than 1.5 nm) remains largely unaffected. This reinforces the concept that Kaps enhance the NPC’s ability to selectively filter macromolecules based on size.

### Transport is a discontinuous process

Kap translocations are predominantly non-directional [67]. Without the Ran-gradient, a net flux of NTR translocations is absent. This is in agreement with our simulations: without active removal of Kaps from the nuclear face of the NPC, almost all Kaps remain inside the CT, moving equally well in both directions. Only when a computational Ran-gradient was introduced, we observed a net movement towards the nuclear side of the pore.

Our simulations show that Kap95 translocation through the NPC occurs in a stepwise manner, characterized by alternating phases of rapid movement and transient pausing (Fig. 4 and Suppl. Movie S2). This behavior closely mirrors recent single-molecule tracking data from human NPCs [21], which revealed similarly intermittent dynamics and a preference for peripheral transport routes. In our simulations, translocating Kaps largely avoid the central axis (*r* ≤ 5 nm) and instead move along the inner surface of the GLFG-ring (Fig. 5a), where the binding avidity is high, i.e., FG-binding sites are abundant and the protein density is relatively low [37, 68].

Kap95 molecules exhibit substantial variation in translocation speed, modulated by their local interactions within the CT. In particular, they slow down when engaging extensively with FG motifs in the GLFG-ring, whereas they move more rapidly along the interface between the GLFG-ring and the Nsp1 percolation zone (Suppl. Movies S2 and S3). This suggests that selective transport is governed not only by the spatial distribution of FG motifs but also by local temporal openings in the protein meshwork, together creating regions of high binding avidity.

Importantly, our results also suggest that there are not two distinct populations of fast and slow Kaps—rather, individual Kaps alternate between more and less mobile phases depending on their local interactions within the CT. All Kaps exhibit dwell phases at key locations (e.g., cytoplasmic face, GLFG-center, nucleoplasmic face), temporarily screening FG-interactions for others. The same Kap can later become highly mobile and rapidly traverse the pore. This dynamic and cooperative behavior increases transport efficiency and supports the view that Kap crowding can enhance rather than hinder translocation, consistent with previous suggestions [26, 50].

Our previous work [69] indicates that both hydrophobic and electrostatic interactions contribute to FG-Nup organization and Kap transport. In particular, the GLFG-ring is not only enriched in FG motifs but also in positively charged residues (Suppl. Fig. S2), which may interact with the negatively charged Kap95, so that electrostatic effects likely contribute alongside hydrophobic interactions.

### Nuclear transport is Kap-centric and Nsp1-mediated

Our simulations provide a comprehensive view of transport through the NPC (Fig. 5c). Inert cargoes primarily diffuse through the center of the pore (*r* ≤ 5 nm), where the extended domains of Nsp1 form a highly-dynamic, percolated meshwork. This Nsp1-rich region supports transient voids that allow small inert particles to translocate without strong interactions. In contrast, Kap95 molecules travel along the surface of the GLFG-ring, avoiding the central channel—consistent with recent experimental observations [21] and the reduction-of-dimensionality model [70], in which transport proceeds along a lower-dimensional surface enriched in transient binding sites. The central Nsp1 network modulates the phase state of the GLFG surface, giving it a dynamic and fluidic nature [65] that suppresses excessive cohesion that could otherwise obstruct selective transport.

Kap-mediated transport is further shaped by crowding effects: early-binding Kaps engage with high-avidity FG regions, particularly with the GLFG-ring, and transiently shield subsequent Kaps from strong interactions, thereby facilitating faster movement through the pore. This redistribution of binding opportunities creates a dynamic, self-regulating response that enhances selectivity. While we do not observe two clearly segregated populations of fast and slow Kaps, the reduced avidity at higher Kap concentrations and consistent translocation paths suggest a collective, Kap-centric modulation of NPC selectivity. Notably, the accumulation of Kaps in the central channel gives rise to an hourglass-shaped density that closely resembles the central “plug” reported in [11, 14, 16] based on cryo-EM reconstructions. Rather than representing a static obstruction, our results suggest that this “plug” is a dynamic ensemble of Kaps and FG-Nups that remains permeable due to the transient and reversible nature of FG-Kap interactions.

Overall, our findings fit within the broader thermodynamic framework of virtual gating [71] and align with the reduction-of-dimensionality [70] and Kap-centric models [4]. Our results corroborate the presence of FG–FG cross-links, but in contrast to the selective phase model [72] we observed these cross-links to be highly transient and not requiring active disruption by Kaps. The most striking insight from our residue-scale simulations is the presence of a high-density GLFG-ring and a central dynamic FG–FG percolated meshwork, primarily mediated by Nsp1, with nanosecond-long cross-links. This creates a dynamic environment for nuclear transport that is both Kap-centric and Nsp1-mediated, and aligns with the reduction-of-dimensionality model.

### Outlook

Our residue-scale simulation framework for nucleocytoplasmic transport can serve as a computational transport assay that allows for a systematic study of a broad range of outstanding questions, from probing the role of variations in pore diameter, the effect of cargo size, surface hydrophobicity and number of Kaps per cargo, to studying complete RanGTP-controlled bidirectional transport cycles of proteins and RNA, providing a pathway towards full mechanistic understanding of selective nuclear transport.

## Methods

### Residue-scale model for intrinsically disordered proteins

All molecular dynamics simulations are performed using the 1BPA-1.1 model for intrinsically disordered proteins [38]. In this implicit-solvent coarse-grained model each amino acid is represented by a single interacting bead, carrying residue-specific physicochemical properties (charge and hydrophobicity) and calibrated to accurately reproduce experimentally determined hydrodynamic radii over a range of yeast FG-Nup segments [31, 38]. The 1BPA model incorporates a sequence-specific backbone stiffness by distinguishing between three groups of amino acids (i.e. glycine, proline and other residues) [30]. Furthermore, cation–pi interactions are included by recalibrating the interaction potentials between cationic (R,K) and aromatic (F,Y,W) residues [34].

All simulations are performed using GROMACS [73] molecular dynamics software (version 2019.6), where the stochastic dynamics integrator operates with a time step of 0.02 ps and inverse friction coefficient *γ*^*−*1^ = 50 ps. Simulations are performed at 300 K and physiological salt concentration of 150 mM by setting the Debye screening constant *κ* = 1.27 nm^*−*1^. By modeling amino acids as single interacting beads and using an implicit solvent model, our simulations significantly reduce the number of degrees of freedom compared to all-atom systems, resulting in a smoothed free-energy landscape and accelerated dynamical processes. The friction coefficient used corresponds to a diffusion coefficient about 500 times larger than experimental measurements [38, 42]. Consequently, we scale the time step by a factor of 500, yielding an effective time step of 10 ps, which we use to report our simulation results.

### Residue-scale model of yeast Kap95

For the residue-scale model of yeast Kap95, we used our previously developed model [37] of the unbound state of Kap95 (PDB ID: 3ND2 [74]). The Kaps are modeled as inert proteins, where the secondary and tertiary structure are maintained using an elastic network. Multivalent interactions with FG-Nups are largely preserved by introducing specific FG–binding site interactions and by using the 1BPA electrostatic and cation–pi interactions. The ten binding sites per Kap95 are accurately determined and refined from a computational study [75]. See Ref. [37] for more details on the CG model and calibration procedure.

Although Kap95 is modeled with 10 nominal FG-binding sites, each site is represented by multiple beads capable of interacting with FG motifs. This allows for transient multivalent contacts where a single site may engage more than one FG motif simultaneously. As a result, the average number of FG contacts per Kap95 molecule can exceed 10. Nevertheless, the structural constraints of Kap95—particularly the buried nature of some binding residues—limit the effective number of simultaneous strong interactions.

### Generation of residue-scale model of the yeast NPC

The structural model of the yeast NPC was derived from an integrative structure of an isolated NPC obtained by Kim *et al*. [11]. The origin of the coordinate system was chosen to coincide with the center of mass of the scaffold structure. Multi-resolution (C_α_ or coarser) PDB structures were then exported for all individual scaffold Nups and FG-Nup domains that comprise the eight-fold symmetric NPC structure. When available, the C_α_-positions were extracted and stored for each Nup, with the exclusion of nuclear basket proteins Mlp1, Mlp2 and Nup2, membrane ring Nups Pom33, Pom34 and Ndc1, and the cytoplasm-facing Gle1 (see Suppl. Tab. S1 for an overview of all modeled scaffold proteins). Missing residues or flexible linker regions between folded domains were positioned via a self-avoiding random walk (see Suppl. Methods), thus completing the scaffold structure to the extent possible. The scaffold structure was fixed in space by freezing all non-linker scaffold residues (i.e. their position was not updated during the simulations). Interactions between scaffold residues and disordered FG domains only comprise excluded volume interactions.

A half-open torus-shaped occlusion was then inserted to mimic the nuclear membrane. The minor and major radii of the torus were set to 13.25 nm and 55.5 nm, respectively (Fig. 1f). This is in accordance with constraints used in the integrative modeling underlying the NPC structure [11]. The sterically inert beads that constitute the torus-shaped membrane were given a diameter of 3.0 nm and only interact with the disordered FG domains by means of volume exclusion. Interactions between the torus-shaped occlusion and the flexible linkers or frozen scaffold beads were excluded.

A self-avoiding pseudo-random walk is used to generate initial configurations of the disordered FG-Nup segments in the yeast NPC. This approach produces physically reasonable, non-overlapping chain conformations and significantly accelerates equilibration compared to starting from fully extended chains. Importantly, we find that the FG-Nup distribution consistently converges to a similar equilibrium structure across simulations (with different Kap concentrations), regardless of the specific initial configuration.

The β-propellers of Nup159 (residues 1–382) were included in the model and attached to the disordered FG domain (residues 382–1116). The structure of the β-propeller was preserved by applying a network of harmonic bonds, where harmonic bonds with spring constant *k* = 8000 kJ mol^*−*1^ nm^*−*2^ were applied to any non-neighboring residues with a separation smaller than 1.2 nm. Other studies on the stoichiometry of NPC components [76] indicated that a second copy of Nup1 is likely present, in addition to the structure of Kim *et al*. adapted here. A second copy of Nup1 was therefore added, with the same anchoring coordinates as the first copy of Nup1 (see Suppl. Methods).

### Simulation procedure

Following the generation of the FG-meshwork, we performed an extensive equilibration simulation of 2.5 × 10^8^ steps (2.5 ms) of the complete NPC (in the absence of Kaps) to allow the FG-meshwork to homogenize before the 2.5 × 10^8^ steps (2.5 ms) production run. For simulations with Kaps, we started with the final state from the initial FG-meshwork equilibration. Kap95 proteins were inserted into the NPC using a custom “growing” pipeline (see Suppl. Methods for more details). After the desired number of Kaps were inserted, we performed another equilibration run of 2.5 × 10^8^ steps (2.5 ms) to allow the NPC–Kap system to reach a new equilibrium state, followed by a 2.5 × 10^8^ steps (2.5 ms) production run. Equilibration was verified by monitoring the time evolution of the FG-Nup and Kap95 density distributions, which stabilized after approximately 2.5 ms (Suppl. Figs. S7 and S8).

To simulate the effect of a Ran-gradient, we implemented a recycling process for Kap95 proteins upon reaching the nuclear side of the NPC. Every 5.0 × 10^5^ steps (5 µs) the simulation is paused and the center-of-mass (COM) position of each Kap is determined. If the *z*-coordinate of a Kap is found to be 20 nm below the center of the NPC scaffold, the Kap is deemed to have reached the nuclear side of the pore (Suppl. Fig. S14). The identified Kap is then removed from the nuclear side of the FG-meshwork and reinserted at the cytoplasmic side (*z*_COM_ = − 60 nm). After checking and recycling all Kaps, the simulation is continued. This procedure is continued until an aggregate simulation trajectory of 1.0 × 10^9^ simulation steps (10 ms) is obtained. For these simulations, equilibration was verified based on the translocation rate over time, which stabilized after around 5 ms (Suppl. Fig. S15).

NPC equilibrium simulations were performed on the Hábrók HPC cluster (University of Groningen) using nodes equipped with two AMD 7763 CPUs (128 cores @ 2.45 GHz). Production runs, ranging from 432,448 beads (no Kap95) to 570,208 beads (160 Kap95 molecules), were conducted on two nodes (256 threads total), achieving a performance of 120–150 µs*/*day. Simulations involving Kap recycling were carried out on the Snellius HPC system (SURF, Netherlands), using nodes with dual AMD Rome 7H12 CPUs (2 × 64 cores @ 2.6 GHz), also using two nodes per run, with performance ranging from 160–170 µs*/*day.

### Mapping of density and displacement data to axi-radial distributions

To calculate the axi-radial density distributions, we first defined an axi-radial coordinate system with the origin at the center-of-mass of the NPC scaffold. For each instantaneous configuration of the FG-meshwork, we calculated the 2D density distribution by averaging in the azimuthal direction on a bin grid (0.5 nm bin size) and correcting for the available volume of each voxel. Finally, we computed the average density distribution over all frames of the equilbrium trajectory. For the simulation snapshots of selected Kaps, the density distributions were calculated over short time windows of 25 µs.

To calculate the spatial distribution of slow and fast Kap displacements, we followed these steps: First, for each Kap, we calculated the center-of-mass displacement vector as the vector from the center-of-mass at time *i* to that at time *i* + window size (window size is 50 µs). Time windows that contained a recycling event were excluded from the analysis. Next, we determined the magnitude of the displacement by taking the vector norm. To correlate these displacements with positions, we determined the average center-of-mass position of each Kap during the selected time window. Using a rolling average, we obtained the displacement along the (average) trajectory of each Kap. The displacements were sorted from smallest to largest, allowing us to identify the slow and fast Kaps by selecting the 25 % smallest and 25 % largest displacements, respectively. For the selected slow and fast displacements, we averaged the associated center-of-mass positions. These positions were then converted from 3D Cartesian coordinates to 2D axi-radial coordinates by averaging over the azimuthal direction, resulting in the number density distribution of slow and fast Kaps.

### Contact and interaction lifetime analysis

To characterize the intermolecular interactions in the NPC, we used the following steps. For each trajectory frame, sampled every 500 ns over the 2.5 ms equilibrium trajectory, we computed a contact matrix using MDAnalysis [77, 78]. This matrix described all interresidue contacts between FG-Nups or between FG-Nups and Kaps. In all analyses only the FG domains and only intermolecular contacts were considered. Residue pairs were considered to be in contact if the distance between their beads was less than 0.7 nm. The contact matrices were then averaged across all trajectory frames to generate the final average contact map.

Distributions of residue–residue contact lifetimes are calculated using a transition-based or core-state approach [79, 80], in which two separate cut-offs define the formation and breaking of a contact between residues. This approach reduces noise from transient (“fly-by”) contacts and more accurately captures true binding events. We have adapted the method to coarse-grained systems by scaling the cut-offs for formation and breaking to *r*_form,CG_ = 0.55 nm and *r*_break,CG_ = 1.2 nm for 1BPA simulations. To verify the robustness of the chosen cut-off values, we compared the contact lifetime distributions for various combinations of cut-off parameters and found no significant difference when changing either of the cut-offs by 10 % (Suppl. Fig. S5).

### Void analysis and PMF barriers

Energy barriers for passive translocation of inert cargo are calculated using the void analysis method described by Winogradoff et al. [42]. The simulation volume was mapped onto a 3D grid, with a grid size of 6 Å. For each configuration of the FG-meshwork, we assessed whether a spherical probe of radius *r*_p_ could be placed at the center of each voxel without overlapping with any amino acid bead of the meshwork or the pore scaffold. The resulting 3D void maps were then converted into 1D potential occupancy maps by calculating the percentage of available voxels in each slice along the pore axis. This potential occupancy function was calculated every 5 × 10^4^ steps (500 ns) of the equilibrium trajectory. The average potential occupancy function over the trajectory was then transformed into an effective PMF curve using Boltzmann inversion. We refer the reader to Ref. [42] for more details on the analysis procedure. The analysis was performed with the codes provided by the paper, with a custom constraint to exclude the volume of the nuclear membrane from the void analysis.

To evaluate the pathway for passive translocation through the CT, we determined the time-averaged distribution of the “voids” within the pore. This was done by computing the 3D void map for each instantaneous configuration of the CT and then averaging these maps over the simulation time. The resulting time-averaged void map represents the probability that each voxel can accommodate an inert protein with radius *r*_p_. Subsequently, this 3D void map was averaged circumferentially around the pore’s axis to produce a 2D void map in (*r, z*) coordinates, as shown in Fig. 4h.

## Data Availability

All data supporting the findings of this study are available within the main text and Supplementary Information. Source data for the main and Supplementary Figures are provided with the paper. Analysis scripts and any additional datasets are available from the corresponding author upon reasonable request.

## Acknowledgements

This work was financially supported by the Netherlands Organization of Scientific Research grant no. OCENW.GROOT.2019.068. This work made use of the Dutch national e-infrastructure with the support of the SURF Cooperative using grant nos. EINF-3563 and EINF-6556. We thank the Center for Information Technology of the University of Groningen for their support and for providing access to the Hábrók high performance computing cluster.

## Notes

### Competing Interest Statement

The authors have declared no competing interest.

### Summary of Updates

Revision of the original manuscript. Added an additional figure and improved the discussion of the results.

