## Supplementary Information for "Kap-centric Nsp1-mediated nuclear transport at full amino-acid resolution"

### Supplementary Methods

#### Generating configurations for the FG-Nup meshwork and flexible linker regions

Coordinates of the disordered FG-domains and certain flexible scaffold segments were not described in full detail in the NPC structure that underlies this study [1]. The starting configurations for FG-domains and certain scaffold segments were therefore generated using an in-house Python code. The placement coordinates for each successive amino acid bead in the amino acid sequence of a disordered FG-Nup segment were assigned using a self-avoiding random walk: starting from the anchoring coordinate, each adjacent residue was inserted based on the location of the previous residue while respecting various constraints. Anchoring coordinates for disordered FG-domains were derived from the position of the closest residue in the FG-Nup's folded anchoring domain (i.e. for virtually all FG-Nups, a folded segment exists in the scaffold region, from which the unfolded FG-domain originates). A folded anchoring region was absent in Nup42, Nup60 and Nup1, hence the protein was anchored based on constraints found by Kim *et al.* [1]. The terminal (anchoring) residues were fixed in space by freezing them, similar to how the scaffold structure was fixed in space.

Starting from the anchoring sites, the procedure for generating FG-Nup configurations can be then described using the following process:

$$\mathbf{x}_{i+1} = \mathbf{x}_i + \mathbf{R}_{\text{dir}} \cdot d_{\text{bond}}, \tag{1}$$

where the coordinate $\mathbf{x}_{i+1}$ of bead $i+1$ is positioned a distance $d_{\text{bond}}$ away from the coordinate $\mathbf{x}_i$ of bead $i$, in the pseudo-random direction given by the unit vector $\mathbf{R}_{\text{dir}}$. To account for the orientation of the FG-Nup (e.g. facing towards the nucleus or cytoplasm) and to prevent dense clusters of amino acid residues near the NPC central channel, a biasing term was included in $\mathbf{R}_{\text{dir}}$:

$$\mathbf{R}_{\text{dir}} = \frac{b\mathbf{R}_{\text{bias}} + (1-b)\mathbf{R}}{|b\mathbf{R}_{\text{bias}} + (1-b)\mathbf{R}|}, \tag{2}$$

where $b$ denotes the biasing strength, $\mathbf{R}_{\text{bias}}$ a unit vector in the bias direction, and $\mathbf{R}$ is a random unit vector. In case of an overlap (distance $\leq \delta$) with a pre-existing particle, a new coordinate was drawn until the overlap is resolved. The overlap distance $\delta$ was chosen in accordance with the the type of particle and segment and decreases with every successive failed placement:

$$\delta_{n+1} = 0.99^n \delta_n, \tag{3}$$

where $n$ indicates the number of trials.

For generating configurations of FG-Nup domains, the value of $d_{\text{bond}}$ was set to the bond length (0.38 nm) and the overlap distance $\delta$ was set to the 1BPA interaction radius of 0.6 nm. A bias strength of 0.2 was used in conjunction with three biasing directions $\mathbf{R}_{\text{bias}}$:

i) Radially inward for the first 100 residues to enable deeply-anchored FG-Nups to protrude out of the scaffold (Nup145N, Nup159, Nup60, Nup1);
ii) Towards a large positive $z$-values for cytoplasm-oriented Nups (Nup100, Nup116, Nup159N, Nup42);
iii) Towards large negative $z$-values for nuclear-oriented Nups (Nup1, Nup60, Nup145N).

After generating configurations for all FG-domains, short molecular dynamics simulations were performed to assess any entanglement of FG-domains. Of all FG-domains, the root mean square fluctuation (RMSF) was calculated for the coordinates $\mathbf{r}_i$:

$$\text{RMSF}_i = \sqrt{\langle(\mathbf{r}_i - \langle\mathbf{r}_i\rangle)^2\rangle}. \tag{4}$$

In case visual inspection or calculation of the RMSF indicated any immobile regions, the procedure was repeated.

The process of generating the configurations of flexible scaffold linker regions comprised an additional constraint. Upon inserting a missing region, the coordinates of the final inserted residue had to be adjacent to the first sequential residue of the scaffold structure. Therefore, a stronger biasing ($b = 0.5$) was used, where the difference vector between the coordinates of the previous residue $\mathbf{x}_i$ and the next known scaffold residue $\mathbf{x}_{\text{final}}$ determined the direction of $\mathbf{R}_{\text{bias}}$. Similarly, the bond length $d_{\text{bond},i+1}$ was enforced based on the number of remaining residues $N - i$ and the distance to $\mathbf{x}_{\text{final}}$:

$$d_{\text{bond},i+1} = \frac{|\mathbf{x}_{\text{final}} - \mathbf{x}_i|}{N - i}, \tag{5}$$

such that the final bead is positioned at the desired location $\mathbf{x}_{\text{final}}$.

#### Placement of Kap95 proteins in the FG-Nup meshwork

To minimally disturb the FG-Nup meshwork when placing Kap95 proteins, we developed a pipeline to grow the Kaps directly in the FG-Nup meshwork. This procedure allows for the placement of a large amount of Kap95 proteins within in a single run.

The coordinates of the Kap95 protein are scaled towards its center of geometry by a factor of 100, effectively reducing it to a single point, but maintaining the rough shape of the protein (within rounding errors). Each shrunken Kap is placed at a randomly assigned location and with a random rotation. The random locations in the NPC are picked in such a way that there is no overlap between the Kaps when they are fully grown, and that there is no overlap between the shrunken Kaps and the FG-Nups or scaffold proteins. Except for volume exclusion interactions, all interactions between Kaps and FG-Nups are switched off during the growing simulations. Furthermore, all Kap95 beads are frozen in place and interactions between and within Kaps are excluded as well. The system is then simulated for a short time ($5 \times 10^4$ steps) to allow FG-Nups to move away from the newly placed beads, as the volume exclusion interactions are still in place. Then, the Kaps are scaled back up such that the radius of the Kap increases by 0.2 nm. The system is again simulated for a short time ($5 \times 10^4$ steps) to let FG-Nup beads move away. This process is repeated until the Kaps have grown back to their original size. Because of rounding errors that occur during the rescaling of the Kap95 coordinates, the shape of the Kap95 proteins is somewhat lost, which can lead to instabilities in later stages of the simulations. Hence, the grown Kap95 coordinates are replaced by the original Kap95 coordinates.

To avoid instabilities during the growing simulations that are caused by the dense packing of Kap95 beads in their scaled-down state, or by the slight overlap between Kap95 beads and FG-Nup beads after each up-scaling iteration, we use a custom interaction potential for the volume exclusion interaction between Kap95 beads and FG-Nups beads. The potential is normalised by multiplying by a scaling factor,

$$f(r_{\text{current}}) = \min\left[1, \left(\frac{r_{\text{current}}}{r_{\text{Kap95}}}\right)^3 + \frac{1}{n_{\text{beads}}}\right],$$

where $r_{\text{current}}$ is the largest distance of any bead in the Kap to its center of geometry in the current simulation and could be considered the Kap's radius, $r_{\text{Kap95}}$ is the radius of the fully grown version of the Kap and $n_{\text{beads}}$ is the number of beads in the Kap ($n_{\text{beads}} = 861$).

| Complex | Protein | Copies/spoke | Structured segments | Unstructured segments | FG domain |
|---|---|---|---|---|---|
| Cytoplasmic-facing | Nup42 | 1 (8) | 383 | 384–430 | 1–382 |
| | Nup116 | 2 (16) | 765, 966–1111 | 766–965, 1112–1113 | 1–764 |
| | Nup100 | 2 (16) | 611, 816–959 | 612–815, 1112–1113 | 1–610 |
| Nup82 complex | Dyn2 | 2 (16) | 7–92 | 1–6 | N/A |
| | Nup82 | 2 (16) | 7–16, 23–120, 123–452, 522–612, 625–669, 678–713 | 1–6, 17–22, 121–122, 453–521, 613–624, 670–677 | N/A |
| | Nup159 | 2 (16) | 1–347, 362–381, 1117–1126, 1211–1239, 1266–1321, 1332–1372, 1382–1412, 1429–1456 | 348–361, 1127–1210, 1240–1265, 1322–1331, 1373–1381, 1413–1428, 1457–1460 | 382–1116 |
| Nup84 complex | Nup84 | 2 (16) | 7–20, 27–80, 96–126, 136–364, 372–483, 506–562, 575–726 | 1–6, 21–26, 81–95, 127–135, 365–371, 484–505, 563–574 | N/A |
| | Nup85 | 2 (16) | 47–126, 132–230, 235–436, 451–492, 496–544, 553–560, 567–585, 590–597, 603–612, 616–634, 638–655, 661–675, 685–699, 707–719, 725–744 | 1–46, 127–131, 231–234, 437–450, 493–495, 545–552, 561–566, 586–589, 598–602, 613–615, 635–637, 656–660, 676–684, 700–706, 720–724 | N/A |
| | Nup120 | 2 (16) | 1–29, 53–305, 311–711, 715–726, 733–746, 754–766, 770–781, 807–818, 821–833, 838–853, 862–879, 884–895, 901–913, 917–931, 942–955, 960–971, 976–987, 994–1008, 1025–1037 | 30–52, 306–310, 712–714, 727–732, 747–753, 767–769, 782–806, 819–820, 834–837, 854–861, 880–883, 896–900, 914–916, 932–941, 956–959, 972–975, 988–993, 1009–1024 | N/A |
| | Nup133 | 2 (16) | 56–78, 86–125, 133–144, 162–184, 193–200, 206–249, 258–480, 490–763, 772–1155 | 1–55, 79–85, 126–132, 145–161, 185–192, 201–205, 250–257, 481–489, 764–771, 1156–1157 | N/A |
| | Nup145C | 2 (16) | 91–99, 126–144, 149–550, 554–560, 566–576, 587–602, 612–624, 631–645, 654–673, 681–689, 703–712 | 1–90, 100–125, 145–148, 551–553, 561–565, 577–586, 603–611, 625–630, 646–653, 674–680, 690–702 | N/A |
| | Seh1 | 2 (16) | 1–248, 288–346 | 249–287, 347–349 | N/A |
| | Sec13 | 2 (16) | 10–158, 166–297 | 1–9, 159–165 | N/A |
| Inner ring | Nup157 | 2 (16) | 88–289, 301–309, 339–457, 481–515, 535–679, 704–730, 744–775, 786–830, 836–892, 900–916, 921–933, 944–1016, 1039–1141, 1155–1391 | 1–87, 290–300, 310–338, 458–480, 516–534, 680–703, 731–743, 776–785, 831–835, 893–899, 917–920, 934–943, 1017–1038, 1142–1154 | N/A |
| | Nup170 | 2 (16) | 98–299, 311–319, 353–471, 505–537, 574–717, 765–791, 831–862, 884–916, 919–930, 936–992, 1000–1016, 1021–1033, 1044–1116, 1141–1191, 1195–1243, 1257–1502 | 1–97, 300–310, 320–352, 472–504, 538–573, 718–764, 792–830, 863–883, 917–918, 931–935, 993–999, 1017–1020, 1034–1043, 1117–1140, 1192–1194, 1244–1256 | N/A |
| | Nup188 | 2 (16) | 12–34, 40–91, 101–123, 131–166, 174–224, 256–282, 288–304, 318–434, 439–479, 493–508, 515–530, 551–577, 584–605, 608–619, 632–785, 793–889, 892–1100, 1119–1133, 1157–1241, 1247–1265, 1276–1292, 1303–1322, 1332–1354, 1383–1567, 1593–1628, 1633–1652 | 1–11, 35–39, 92–100, 124–130, 167–173, 225–255, 283–287, 305–317, 435–438, 480–492, 509–514, 531–550, 578–583, 606–607, 620–631, 786–792, 890–891, 1101–1118, 1134–1156, 1242–1246, 1266–1275, 1293–1302, 1323–1331, 1355–1382, 1568–1592, 1629–1632, 1653–1655 | N/A |
| | Nup192 | 2 (16) | 1–362, 417–574, 602–798, 814–849, 857–953, 961–1126, 1137–1226, 1234–1258, 1272–1366, 1371–1418, 1421–1502, 1511–1559, 1584–1590, 1597–1619, 1623–1644, 1651–1683 | 363–416, 575–601, 799–813, 850–856, 954–960, 1127–1136, 1227–1233, 1259–1271, 1367–1370, 1419–1420, 1503–1510, 1560–1583, 1591–1596, 1620–1622, 1645–1650 | N/A |
| Membrane ring | Nup53 | 2 (16) | 248–284, 304–360 | 1–247, 285–303, 361–475 | N/A |
| | Nup59 | 2 (16) | 266–302, 346–402 | 1–265, 303–345, 403–528 | N/A |
| | Pom152 | 2 (16) | | | N/A |
| Nic96 complex | Nic96 | 4 (32) | 20–56, 205–360, 366–374, 405–444, 455–515, 533–747, 753–835 | 1–19, 57–204, 361–365, 375–404, 445–454, 516–532, 748–752, 836–839 | N/A |
| | Nup49 | 4 (32) | 270–359, 369–407, 433–472 | 360–368, 408–432 | 1–269 |
| | Nup57 | 4 (32) | 287–423, 433–476, 505–541 | 424–432, 477–504 | 1–286 |
| | Nsp1* | 6 (48) | 637–727, 742–778, 788–823 | 728–741, 779-787 | 1–636 |
| Nucleus-facing | Nup1 | 2 (16)** | 1, 351 | 2–350 | 352–1076 |
| | Nup60 | 2 (16) | 1, 398 | 2–397 | 399–539 |
| | Nup145N | 2 (16) | 243, 459–605 | 244–458 | 1–242 |

**Tab. S1.** Overview of stoichiometry and modeled domains (folded or unstructured) in a 1BPA model of the yeast NPC. Scaffold Nups that were missing from the integrative structure or did not comprise any structured domains (e.g. Mlp1, Mlp2, Pom33, Pom34, Ndc1, Nup2) were not included in the model, and are excluded from this overview.

| FG-Nup | Kim *et al.* [1] | Ghavami *et al.* [2] | Huang *et al.* [3] | Winogradoff *et al.* (model 1) [4] | Winogradoff *et al.* (model 2) [4] | Current study |
|---|---|---|---|---|---|---|
| Nup159 | 482–1116 | 390–1460 | 388–1082 | 1–760 | 382–1141 | 382–1116 |
| Nup42 | 1–363 | 1–430 | 1–382 | N/A | 1–382 | 1–382 |
| Nup116 | 1–750 | 1–726 | 1–966 | 1–760 | 1–760 | 1–764 |
| Nup100 | 1–550 | 1–816 | 1–800 | 1–560 | 1–560 | 1–610 |
| Nsp1 | 1–636 | 1–620 | 1–601 | 1–620 | 1–620 | 1–636 |
| Nup49 | 1–269 | 1–472 | 1–270 | 1–220 | 1–220 | 1–269 |
| Nup57 | 1–286 | 1–541 | 1–287 | 1–220 | 1–220 | 1–286 |
| Nup145N | 1–200 | 1–896 | 1–426 | 1–220 | 1–220 | 1–242 |
| Nup1 | 352–1049 | 1–934 | 201–1076 | 1–720 | 357–1076 | 352–1076 |
| Nup60 | 399–498 | 1–539 | 351–539 | 1–120 | 420–539 | 399–539 |
| Nup2 | N/A | N/A | 1–720 | N/A | 1–720 | N/A |

**Tab. S2.** Comparison of FG-Nup domain lengths, when high-charged FG-Nup domains are considered parts of the NPC scaffold (i.e. flexible linkers) or parts of the FG domain. The FG domain lengths considered in various recent computational studies are included as a reference.

### Supplementary Figures

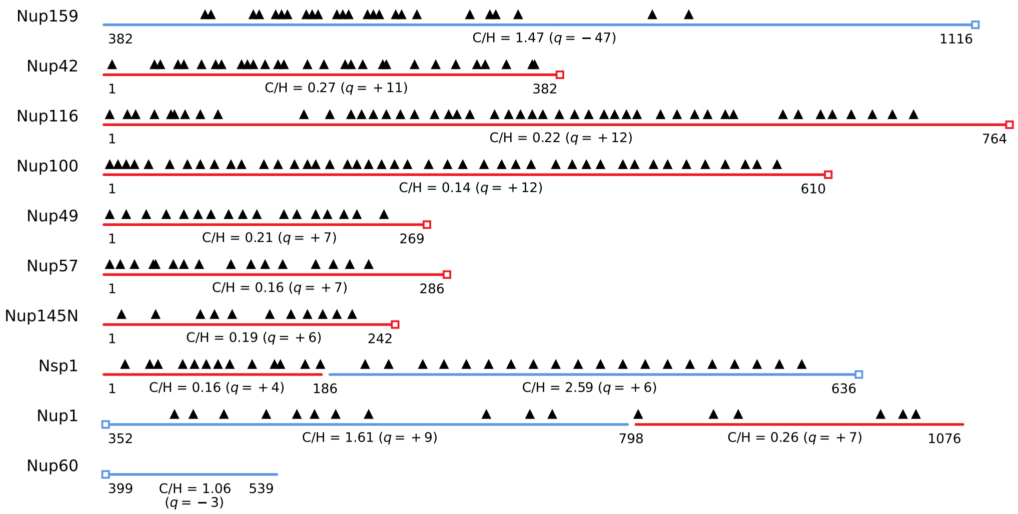

**Fig. S1. FG-Nup domains in the NPC.** Black triangles indicate the positions of FG-motifs, while squares mark the terminal ends anchored to the NPC scaffold. Domains are colored red (cohesive) and blue (noncohesive), following the classification by Yamada *et al.* [5]. The charge-to-hydrophobicity ratio (C/H) is calculated as the number of charged residues (R, K, D, E) divided by the number of hydrophobic residues (F, Y, W, I, L, V). The net charge of each domain, denoted as $q$, is also reported. For reference, Kap95 has a C/H ratio of 0.63 and a net charge of $q = -45$.

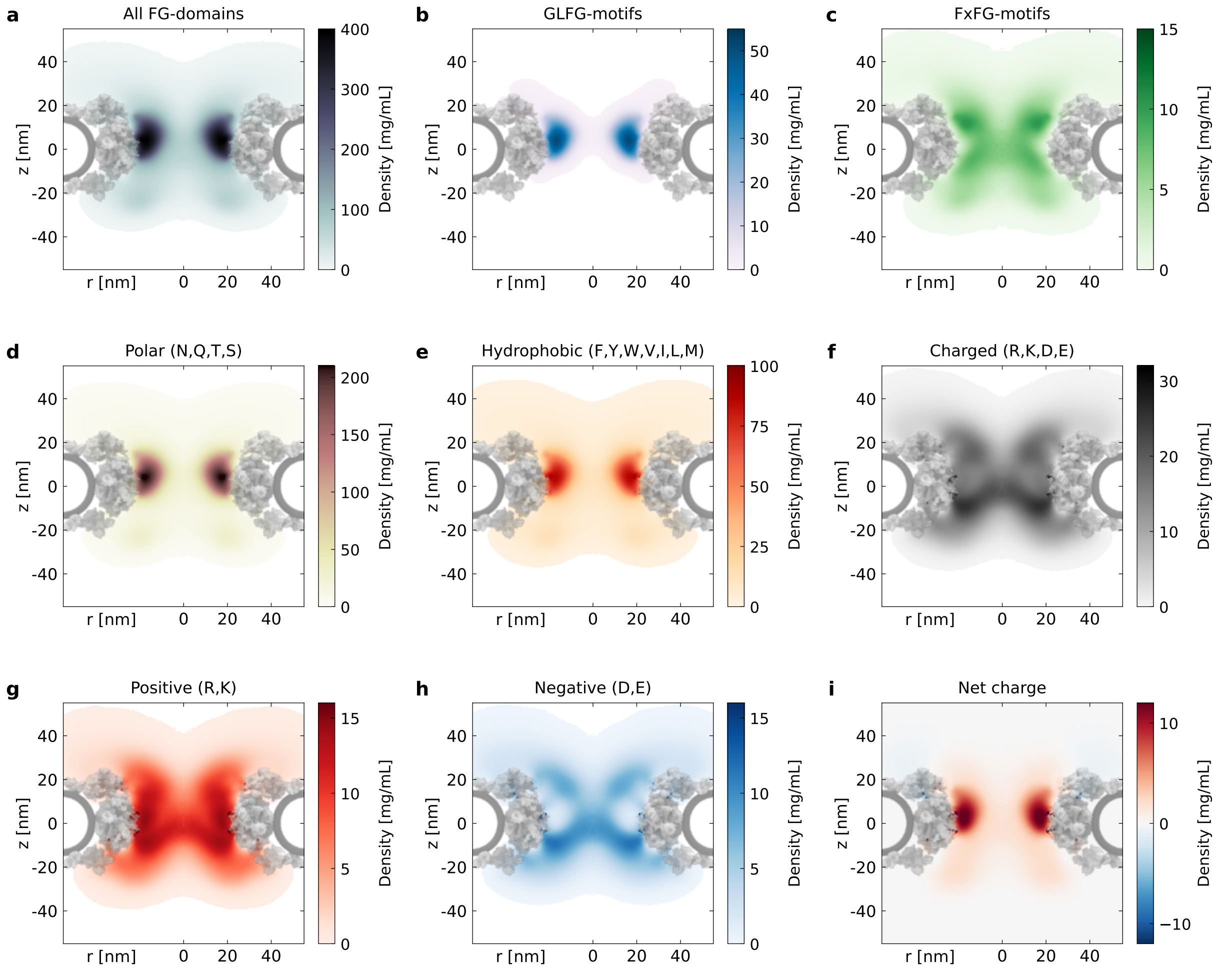

**Fig. S2. Axi-radial, azimuthally and time-averaged density maps of FG-Nups and various types of residues.** (**a**) Total density of FG-domains. A dense ring is formed in the central channel of the NPC. (**b**) GLFG-motifs localize predominantly in the central ring. (**c**) FxFG-motifs, primarily present in Nsp1, localize in the central channel of the NPC. (**d,e**) Hydrophobic and polar residues play a major role in the formation of dense regions in the NPC. (**f–i**) Distribution of charged residues in the NPC. The largest charge density exists in the central channel, at the nuclear side of the GLFG-ring. A segregation of positively and negatively-charged residues (panels g,h) exists and is most pronounced in the central ring. This same effect also comes forward in the accumulation of positive and negative net charge regions (panel i).

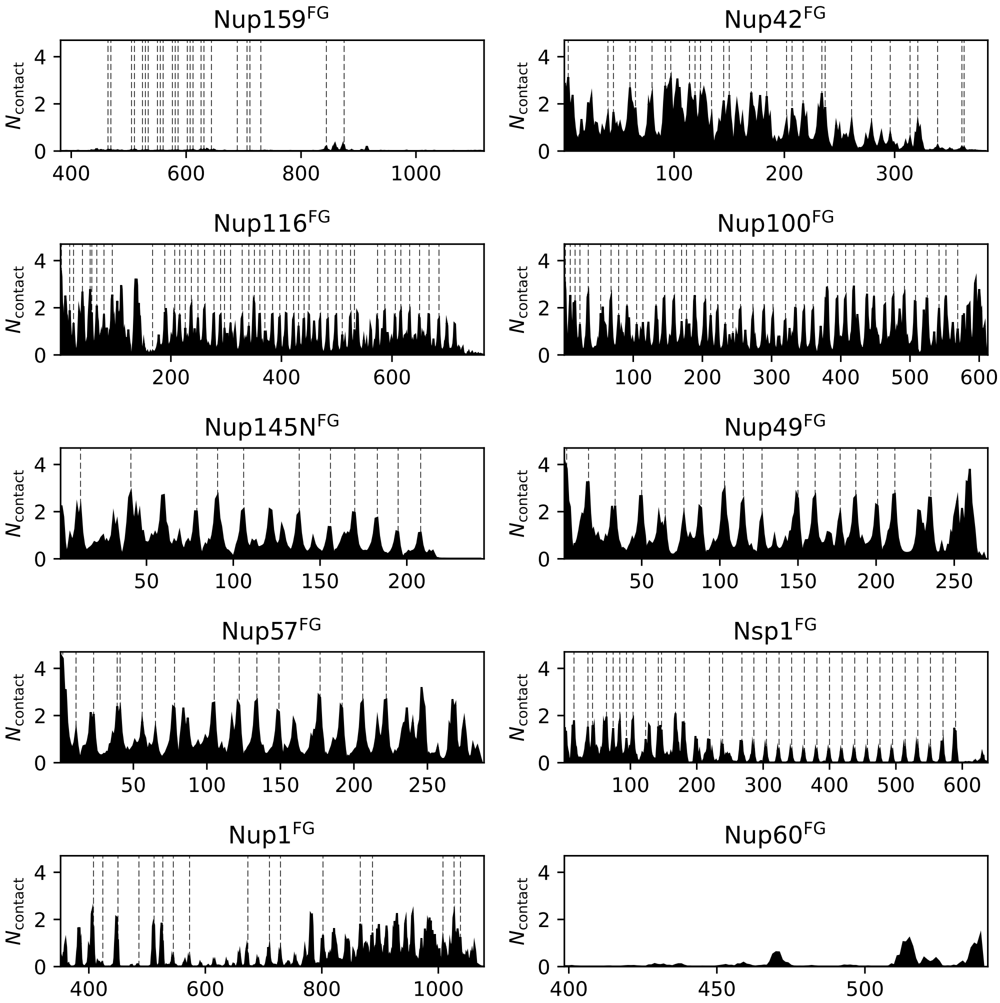

**Fig. S3. Intermolecular interactions in the NPC.** Time-averaged number of intermolecular contacts per protein replica as a function of residue number. The dashed lines indicate the location of FG-motifs. The contact maps show high similarity with those of the FG-Nup condensates (Fig. 3 in [6]) and to the contact maps of single chain simulations (Fig. S12 in [6]).

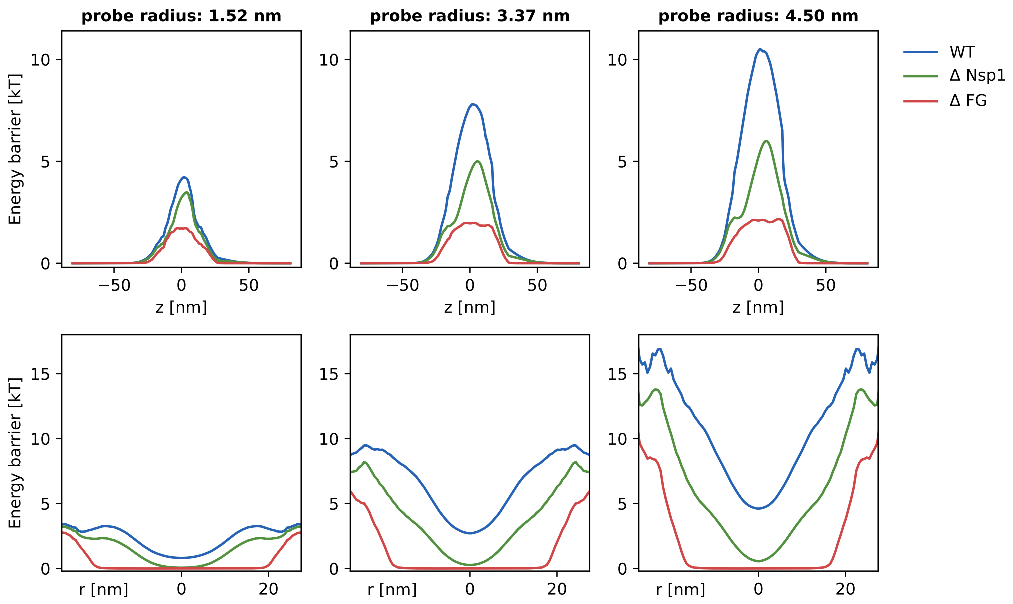

**Fig. S4. The energy barrier for passive translocation in the NPC without Kaps.** The energy barrier for passive translocation (**top**) in the *z*-direction along the pore axis and (**bottom**) in the *r*-direction at the center of the pore for the full NPC (blue), when Nsp1 is removed (green) and when all FG domains are removed (red). Note that the analysis considers the same simulation trajectory in all three cases; the Nup deletions are done during post-processing.

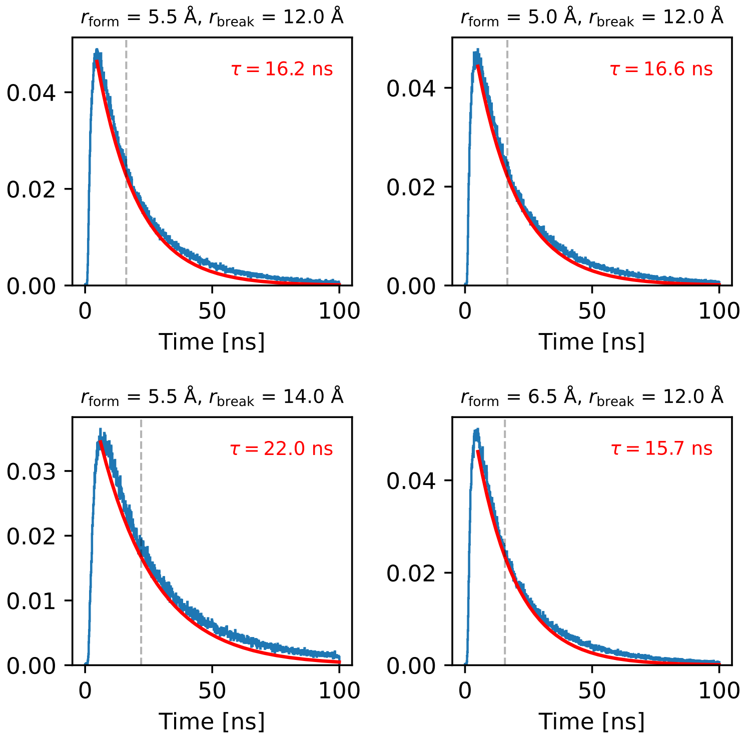

**Fig. S5. Distribution of the lifetimes of FG–FG contacts in the NPC (without Kaps).** The probability density of observed lifetimes is shown in blue, and the red curve represents the best-fit mono-exponential function: $P(t) = (1/\tau)\exp(-t/\tau)$, where $\tau$ is the characteristic lifetime. The fitting is performed from the time corresponding to the peak of the distribution, to avoid including transient "fly-by" contacts. The figures show the contact lifetime distribution for different choices of cut-off distances for the formation and breaking of a contact. The characteristic lifetime $\tau$ extracted from the fit is reported in the figure and marked by a dashed gray line.

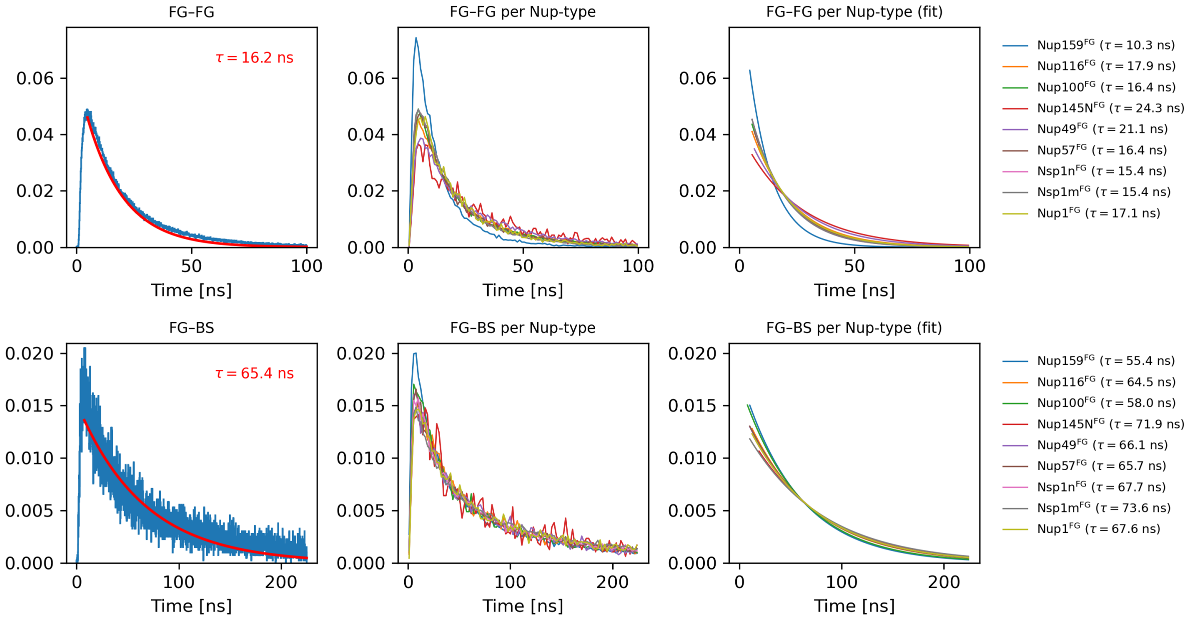

**Fig. S6. Interaction lifetime distributions for FG–FG and FG–BS contacts. Top panels**: FG–FG contact lifetimes. (**Left**) Lifetime distribution for all FG–FG interactions (blue), overlaid with a mono-exponential fit (red), yielding the characteristic lifetime $\tau$. (**Middle**) Lifetime distributions separated by FG-Nup type. (**Right**) Corresponding mono-exponential fits for each FG-Nup type; characteristic lifetimes are indicated in the legend. **Bottom panels**: FG–BS (FG-Nup to Kap95 binding site) contact lifetimes, shown in the same layout as the top row.

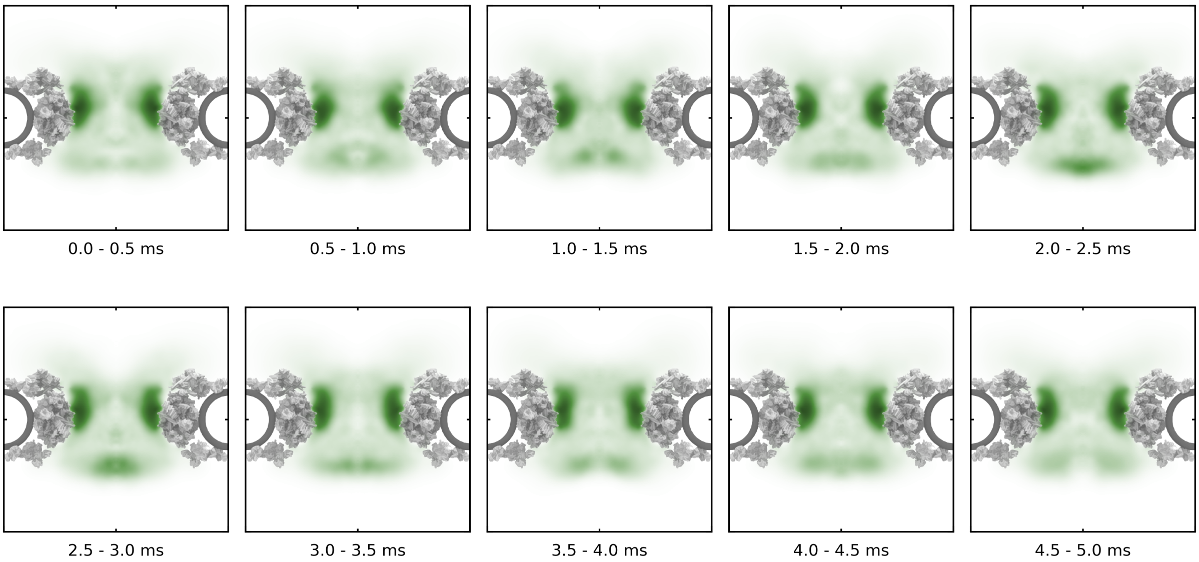

**Fig. S7. Equilibrium verification of NPC with 80 Kaps simulation from Nup density distribution.**

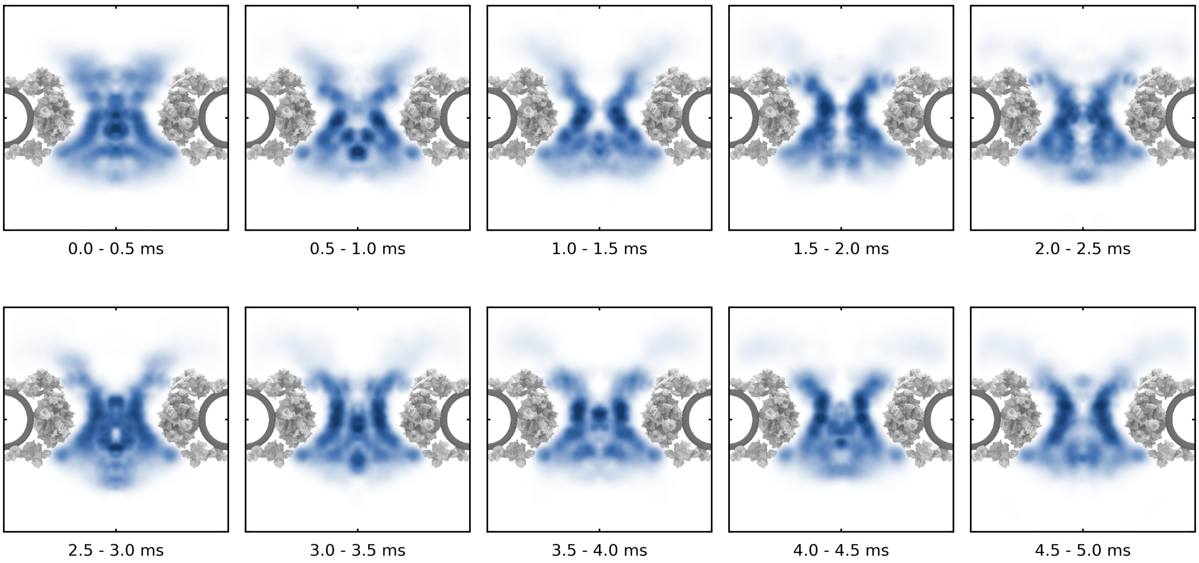

**Fig. S8. Equilibrium verification of NPC with 80 Kaps simulation from Kap95 density distribution.**

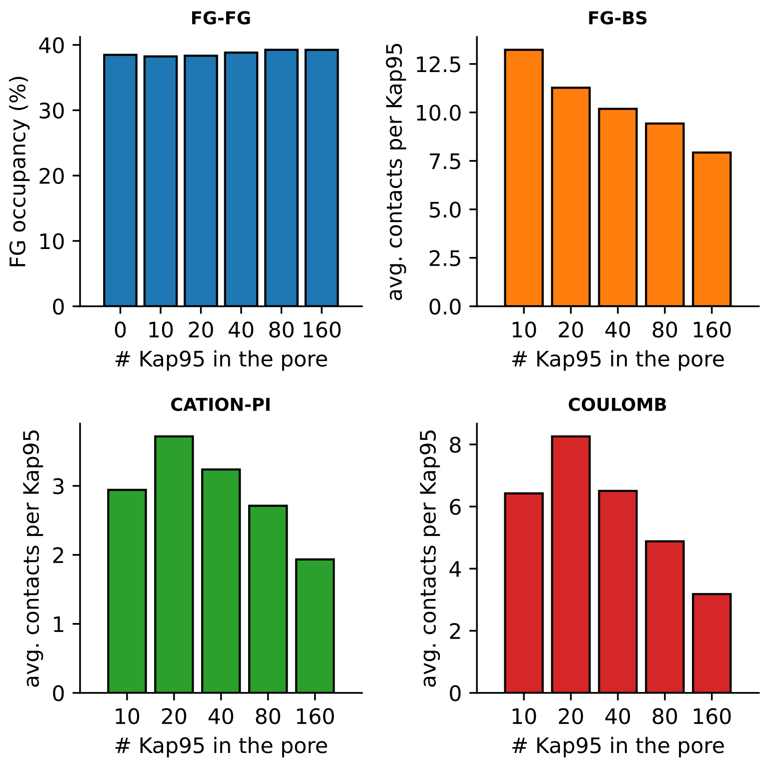

**Fig. S9. Contact statistics in the NPC.** The FG–FG occupancy (i.e. the fraction of FG-motifs that is cross-linked with another FG-motif) is independent of the number of Kap95 proteins in the pore. The average number of FG-interactions per Kap decreases as the number of Kap95 proteins in the pore increases. These two observations combined suggest that the Kaps compete with each other for the available FG-motifs, rather than competing with the FG-meshwork.

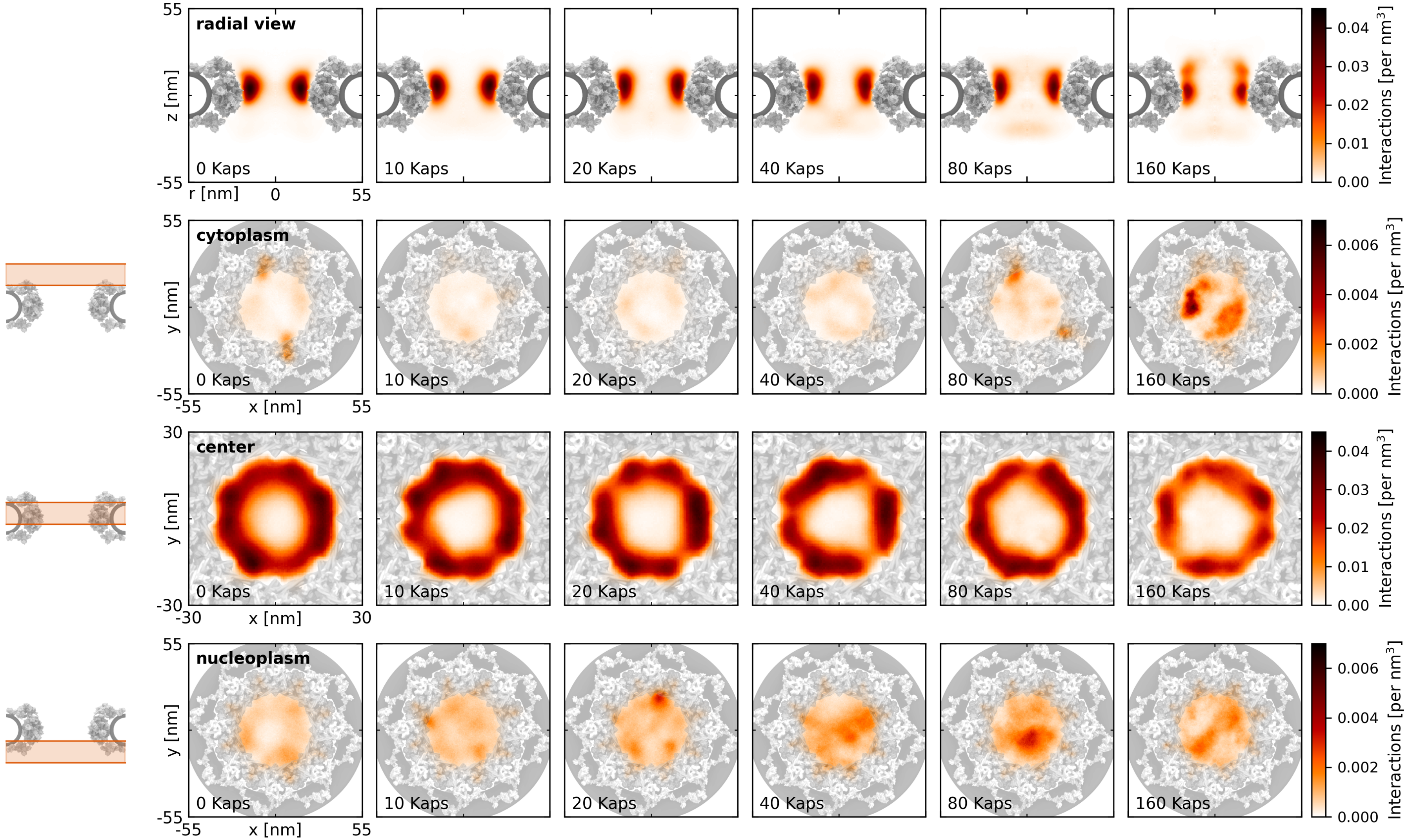

**Fig. S10. Distribution of FG–FG interactions in the pore at varying Kap95 concentrations.** Cross-sections are taken over three axial slices corresponding to different pore regions: 20 to 40 nm (cytoplasmic side), −5 to 15 nm (central region, approximately where the GLFG-ring is located), and −30 to −10 nm (nucleoplasmic side).

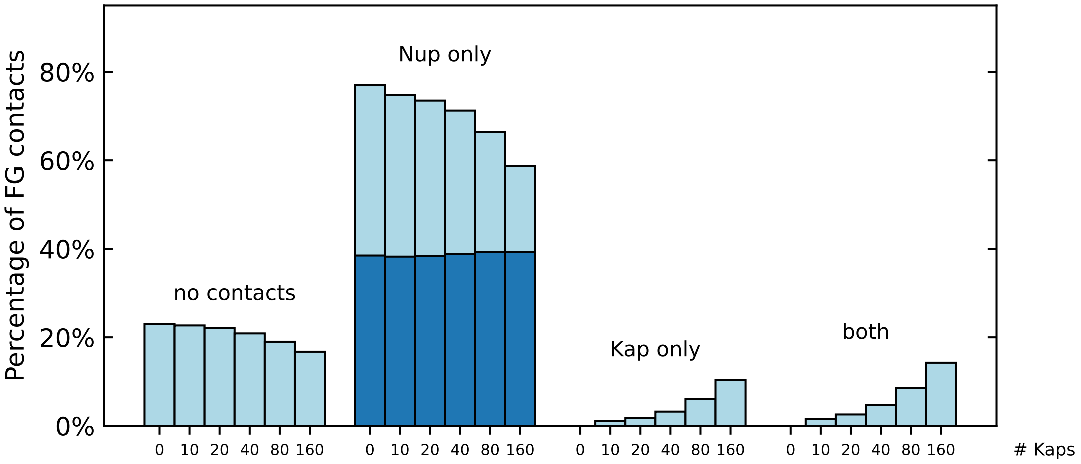

**Fig. S11. Distribution of FG-motif contacts at varying Kap95 concentrations.** Bar plots showing the percentage of FG motifs falling into four categories: unbound, interacting only with FG-Nups, only with Kap95, or simultaneously with both (due to the coarse-grained representation, multiple contacts per FG motif are possible). The dark blue overlay indicates the percentage of FG motifs specifically engaged in FG–FG interactions, cf. Suppl. Fig. S9.

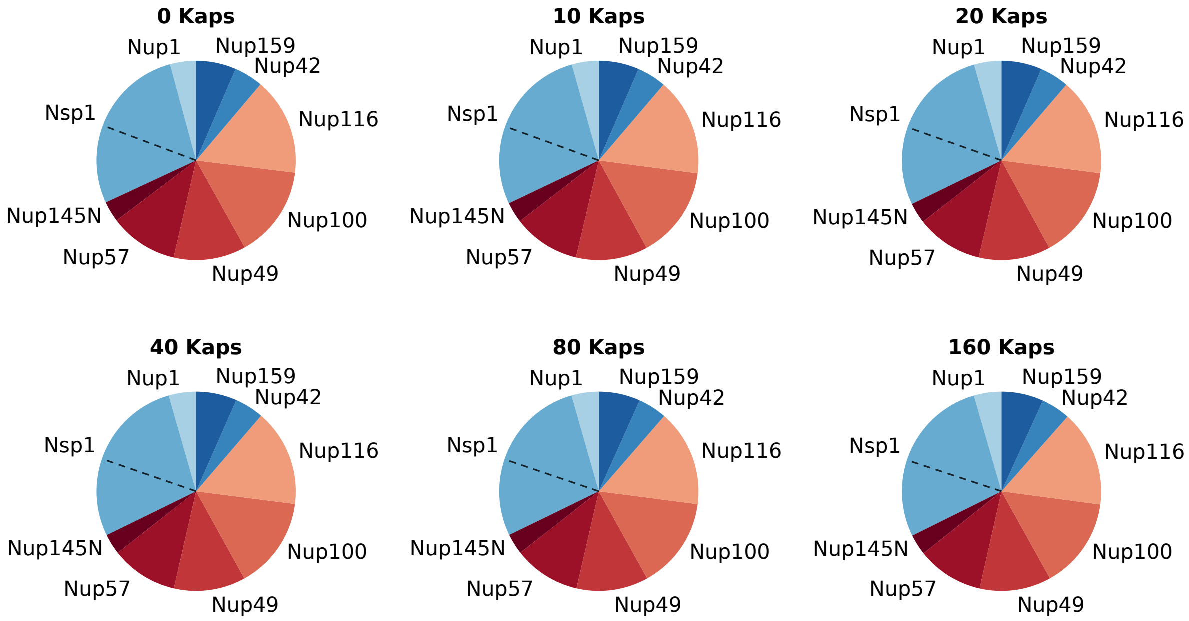

**Fig. S12. Relative fractions of FG–FG interactions by Nup type across Kap95 concentrations.** Pie charts display the distribution of FG–FG contacts among different FG-Nup types at varying Kap95 concentrations. The composition is remarkably stable across conditions, indicating that FG–FG interactions are largely insensitive to Kap loading. GLFG-rich Nups (red) contribute the most to FG–FG contacts, but substantial fractions are also formed by FxFG-type Nups and Nsp1 (blue). Dashed lines within the Nsp1 segment distinguish between contacts formed by its collapsed and extended domains.

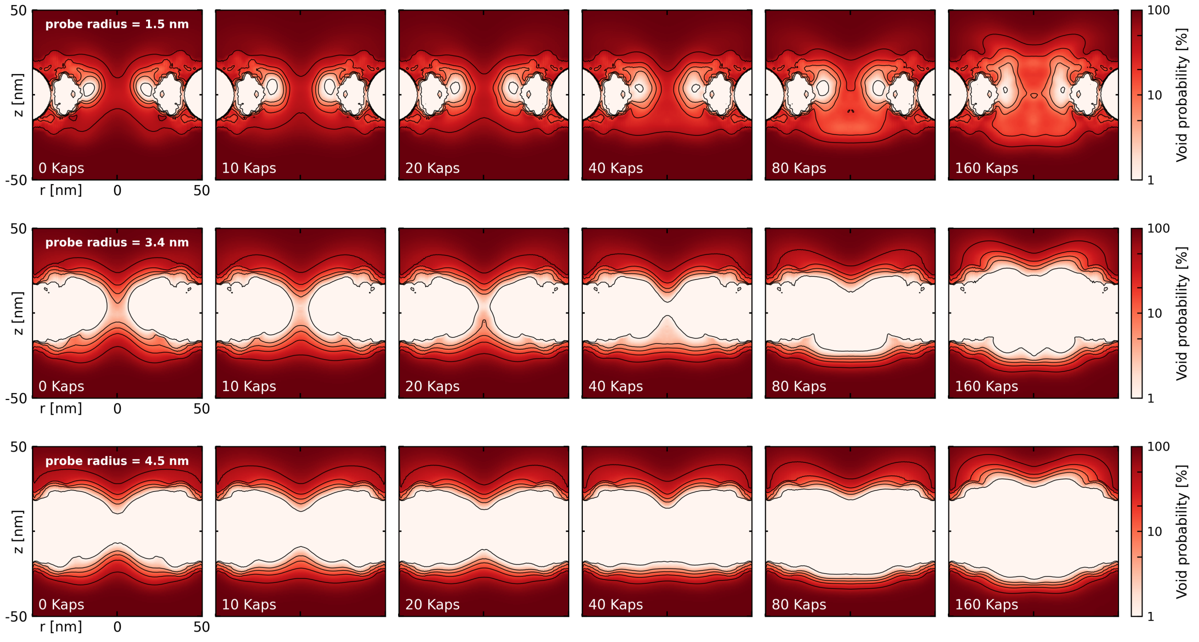

**Fig. S13. Energy barrier for passive translocation at varying Kap concentrations.** Radially-averaged void distribution in the NPC at varying concentrations of Kap95, calculated for a probe radius of 1.5 nm (e.g. ubiquitin), 3.4 nm (e.g. BSA) and 4.5 nm. Contour lines are added at 1, 5, 10, 20 and 50. A clear difference is observed between small inert cargoes (radius = 1.5 nm) and large inert cargos (radius $\geq$ 3.4 nm): For small cargoes there is a clear transport channel through the center of the pore, while for larger inert particles this channel is largely absent. Only for NPCs with a low concentration of Kap95 there is still a transport channel visible.

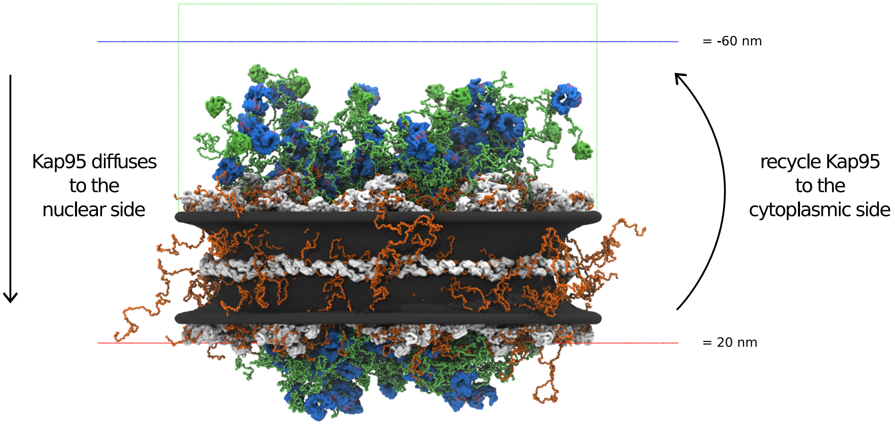

**Fig. S14. Kap95 recycling procedure.** The center-of-mass (COM) position of each Kap is calculated. If the $z$-coordinate of a Kap is 20 nm below the origin of the NPC scaffold (red line), the Kap gets recycled to the cytoplasmic side ($z_{\mathrm{COM}} = -60$ nm, blue line). The Kaps are always contained within a volume near the pore (green box).

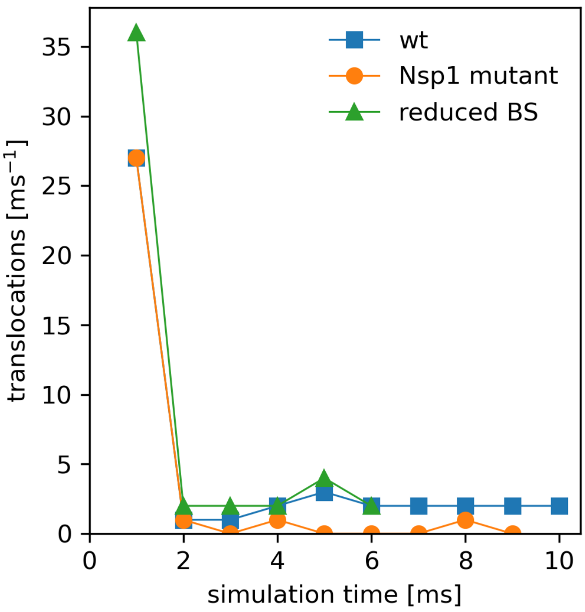

**Fig. S15. Translocation rates in Kap95 recycling simulations.** The translocation rate is measured as the number of Kap95 molecules exiting the pore per 1 ms window. The wild-type (wt) simulation (blue line) shows an initial decline followed by a stable translocation rate of ∼2 Kaps/ms. In a mutant where the FG-binding interactions of the extended domain of Nsp1 (AA 187–636) are turned off (orange line), translocation drops sharply and remains near zero, highlighting the essential role of FG-interactions on the extended domain of Nsp1. In a second mutant, where only half of the Kap95 FG-binding sites (odd-numbered sites) are retained (green line), the translocation rate initially declines but eventually stabilizes at a level comparable to wt, indicating that a reduced number of binding sites can still support effective transport.

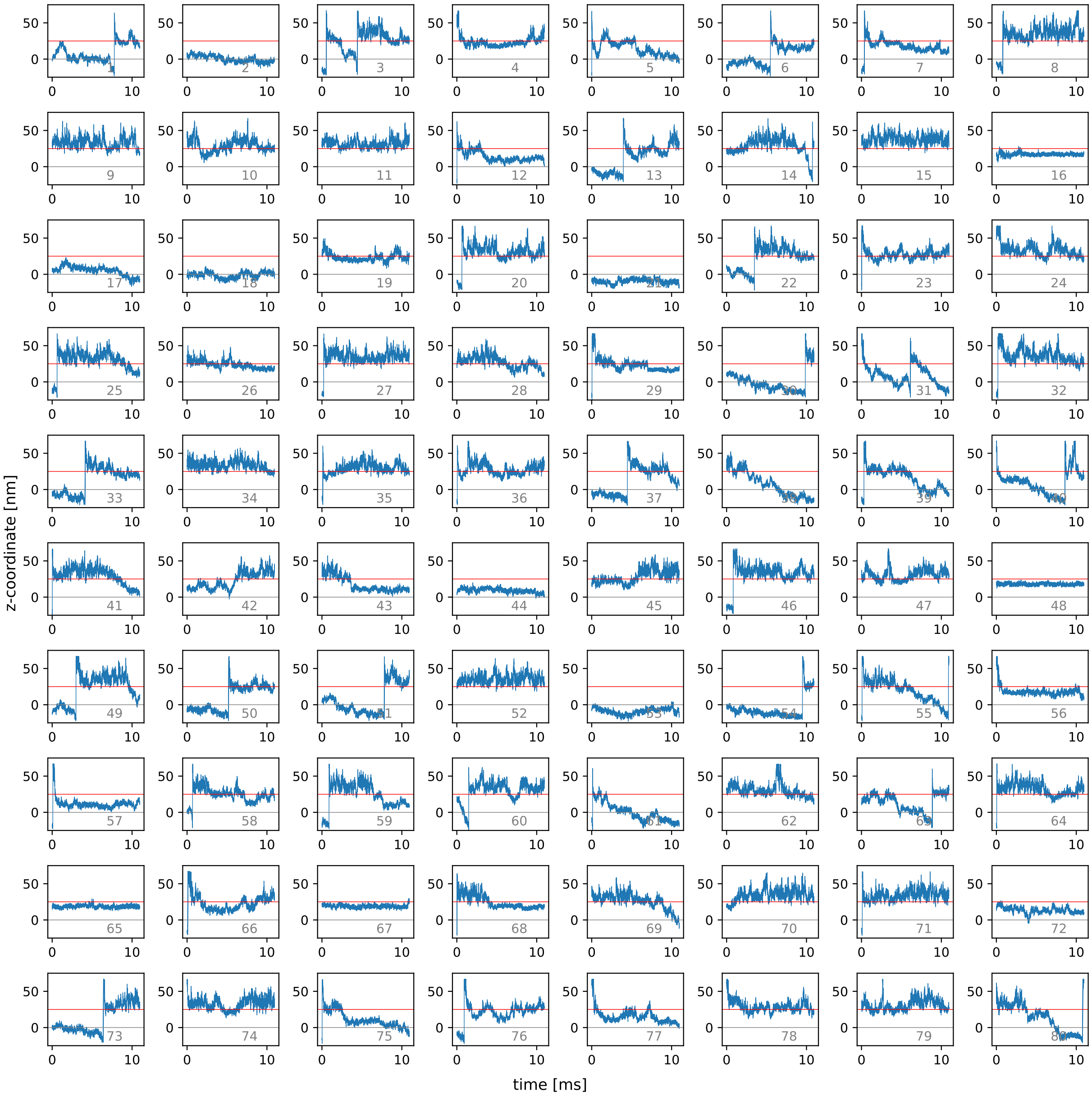

**Fig. S16. Axial trajectories of individual Kap95 molecules during the recycling simulation.** Each subplot displays the $z$-coordinate (along the pore axis) over time for one of the 80 Kap95 molecules in the simulation. The red horizontal line at $z = 25\,\text{nm}$ marks the top of the NPC scaffold on the cytoplasmic side, while the grey line at $z = 0\,\text{nm}$ marks the center of the pore.

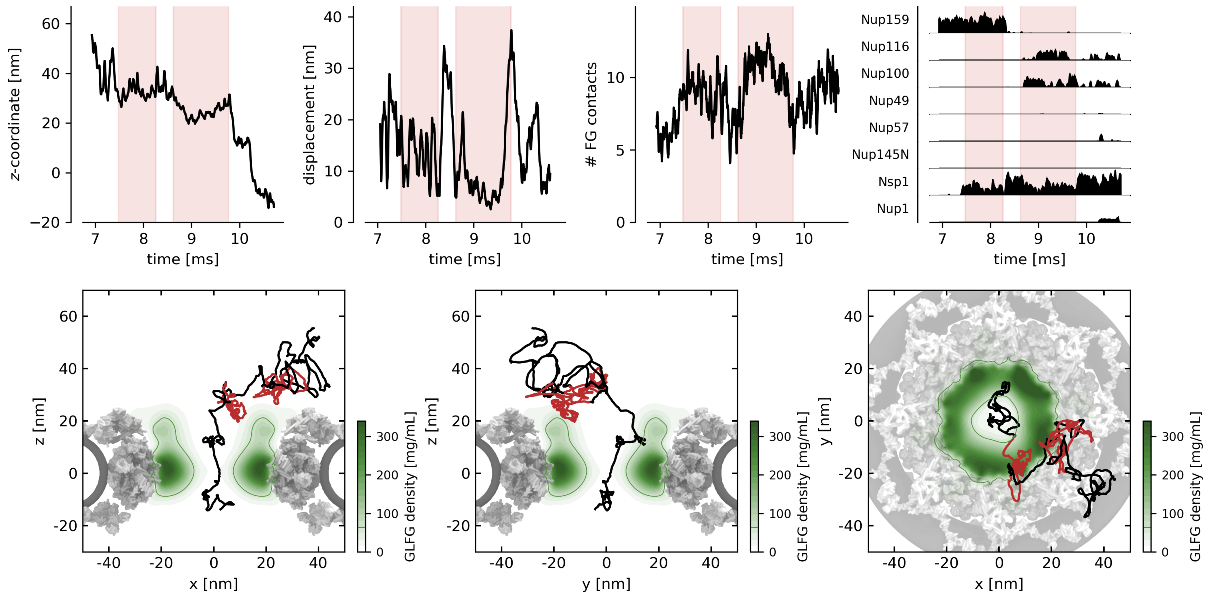

**Fig. S17. Detailed analysis of a single Kap95 translocation event (KAP #14)**. Top panels (left to right): (1) Axial position ($z$-coordinate) of the Kap95 as a function of time, showing its progression through the pore. (2) Instantaneous speed of the Kap, calculated as the displacement over 50 µs time windows. (3) Number of concurrent FG-motif interactions over time. (4) Breakdown of FG-contacts per FG-Nup type. In all four top panels, red intervals indicate dwell times within the FG-meshwork. Bottom panels (left to right): (1) 2D $x$–$z$ projection of the Kap trajectory, overlaid on the average density map of GLFG-Nups. (2) Same trajectory shown in the $y$–$z$ plane. (3) Same trajectory shown in the $x$–$y$ plane (top view of the pore). The red segments in the trajectory correspond to the time intervals marked in red in the top panels.

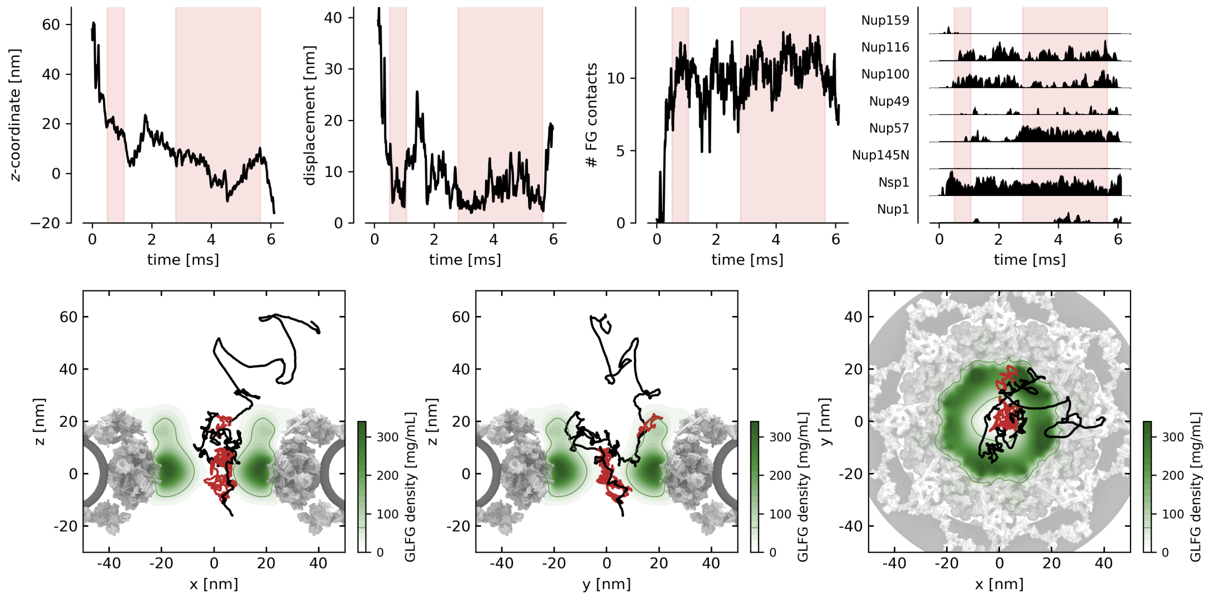

**Fig. S18. Detailed analysis of a single Kap95 translocation event (KAP #31)**. Top panels (left to right): (1) Axial position ($z$-coordinate) of the Kap95 as a function of time, showing its progression through the pore. (2) Instantaneous speed of the Kap, calculated as the displacement over 50 µs time windows. (3) Number of concurrent FG-motif interactions over time. (4) Breakdown of FG-contacts per FG-Nup type. In all four top panels, red intervals indicate dwell times within the FG-meshwork. Bottom panels (left to right): (1) 2D $x$–$z$ projection of the Kap trajectory, overlaid on the average density map of GLFG-Nups. (2) Same trajectory shown in the $y$–$z$ plane. (3) Same trajectory shown in the $x$–$y$ plane (top view of the pore). The red segments in the trajectory correspond to the time intervals marked in red in the top panels.

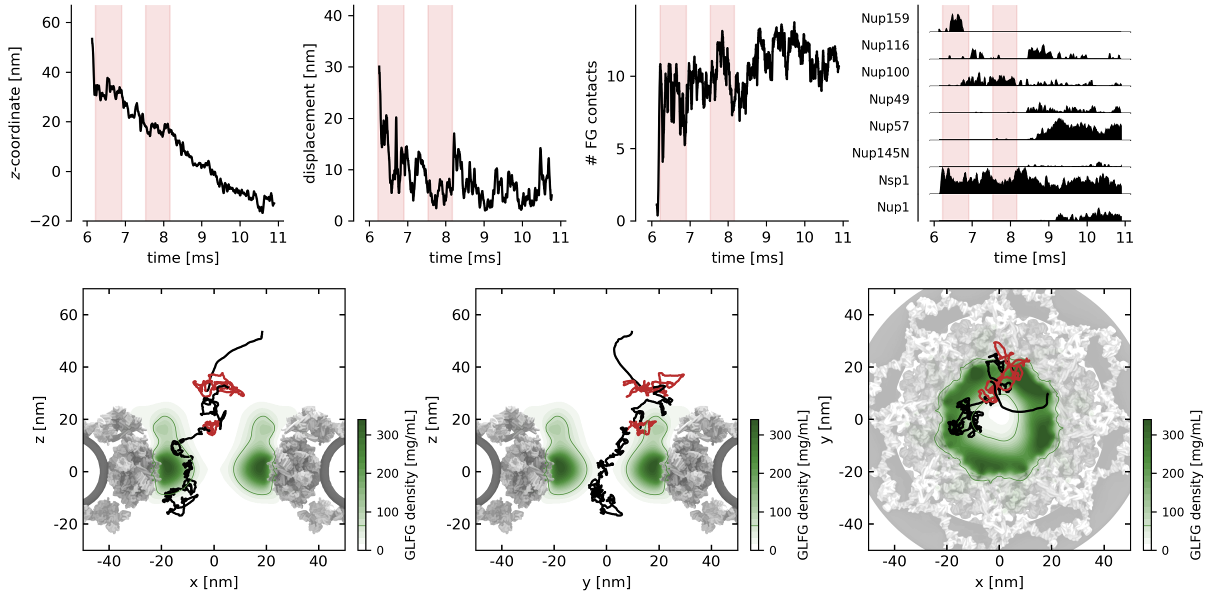

**Fig. S19. Detailed analysis of a single Kap95 translocation event (KAP #31)**. Top panels (left to right): (1) Axial position ($z$-coordinate) of the Kap95 as a function of time, showing its progression through the pore. (2) Instantaneous speed of the Kap, calculated as the displacement over 50 µs time windows. (3) Number of concurrent FG-motif interactions over time. (4) Breakdown of FG-contacts per FG-Nup type. In all four top panels, red intervals indicate dwell times within the FG-meshwork. Bottom panels (left to right): (1) 2D $x$–$z$ projection of the Kap trajectory, overlaid on the average density map of GLFG-Nups. (2) Same trajectory shown in the $y$–$z$ plane. (3) Same trajectory shown in the $x$–$y$ plane (top view of the pore). The red segments in the trajectory correspond to the time intervals marked in red in the top panels.

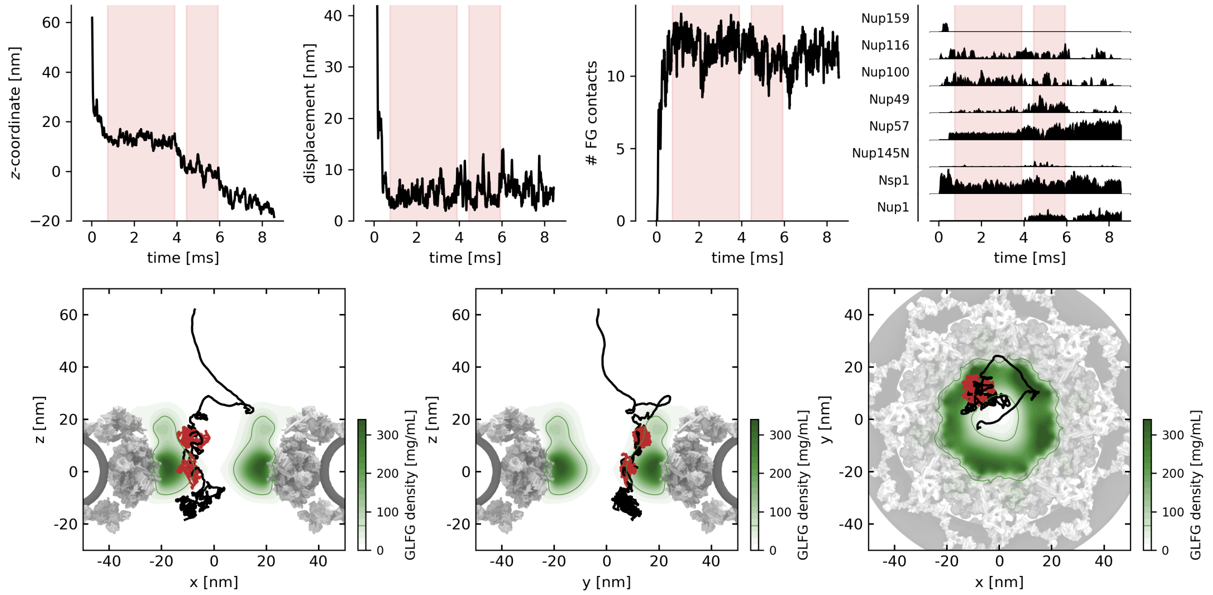

**Fig. S20. Detailed analysis of a single Kap95 translocation event (KAP #40)**. Top panels (left to right): (1) Axial position ($z$-coordinate) of the Kap95 as a function of time, showing its progression through the pore. (2) Instantaneous speed of the Kap, calculated as the displacement over 50 µs time windows. (3) Number of concurrent FG-motif interactions over time. (4) Breakdown of FG-contacts per FG-Nup type. In all four top panels, red intervals indicate dwell times within the FG-meshwork. Bottom panels (left to right): (1) 2D $x$–$z$ projection of the Kap trajectory, overlaid on the average density map of GLFG-Nups. (2) Same trajectory shown in the $y$–$z$ plane. (3) Same trajectory shown in the $x$–$y$ plane (top view of the pore). The red segments in the trajectory correspond to the time intervals marked in red in the top panels.

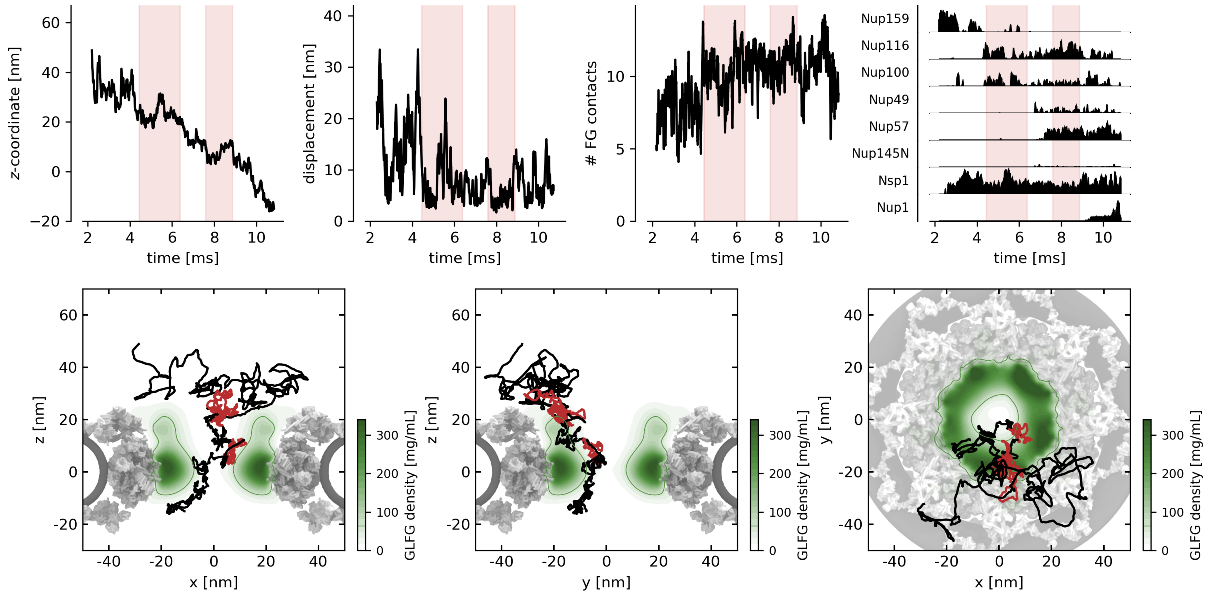

**Fig. S21. Detailed analysis of a single Kap95 translocation event (KAP #55)**. Top panels (left to right): (1) Axial position (*z*-coordinate) of the Kap95 as a function of time, showing its progression through the pore. (2) Instantaneous speed of the Kap, calculated as the displacement over 50 µs time windows. (3) Number of concurrent FG-motif interactions over time. (4) Breakdown of FG-contacts per FG-Nup type. In all four top panels, red intervals indicate dwell times within the FG-meshwork. Bottom panels (left to right): (1) 2D *x*–*z* projection of the Kap trajectory, overlaid on the average density map of GLFG-Nups. (2) Same trajectory shown in the *y*–*z* plane. (3) Same trajectory shown in the *x*–*y* plane (top view of the pore). The red segments in the trajectory correspond to the time intervals marked in red in the top panels.

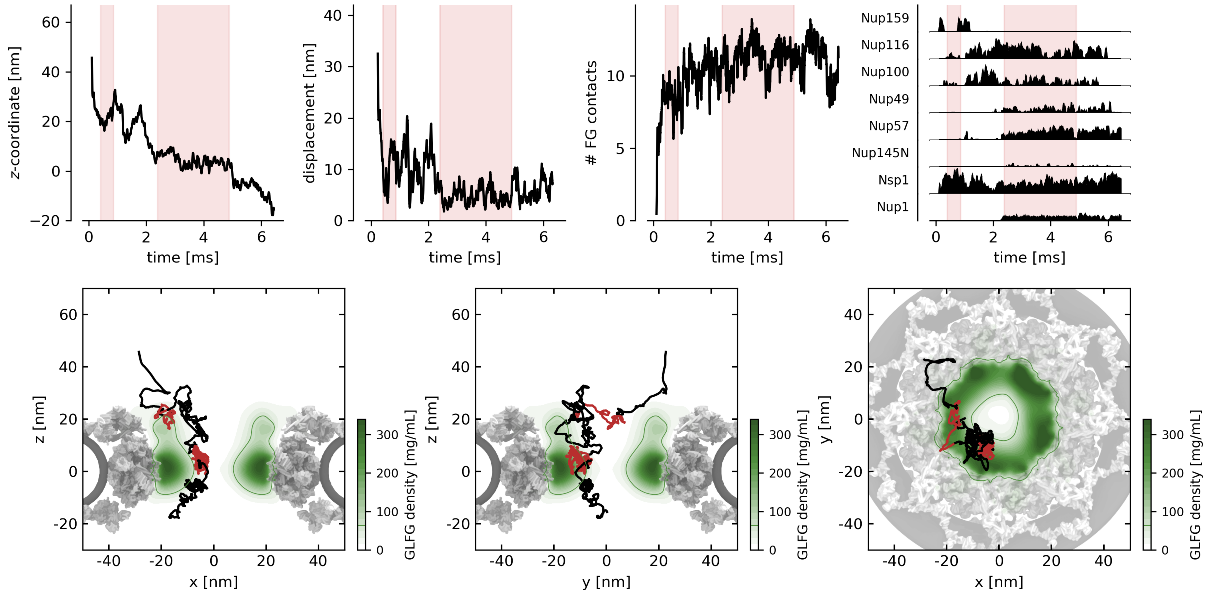

**Fig. S22. Detailed analysis of a single Kap95 translocation event (KAP #61)**. Top panels (left to right): (1) Axial position (*z*-coordinate) of the Kap95 as a function of time, showing its progression through the pore. (2) Instantaneous speed of the Kap, calculated as the displacement over 50 µs time windows. (3) Number of concurrent FG-motif interactions over time. (4) Breakdown of FG-contacts per FG-Nup type. In all four top panels, red intervals indicate dwell times within the FG-meshwork. Bottom panels (left to right): (1) 2D *x*–*z* projection of the Kap trajectory, overlaid on the average density map of GLFG-Nups. (2) Same trajectory shown in the *y*–*z* plane. (3) Same trajectory shown in the *x*–*y* plane (top view of the pore). The red segments in the trajectory correspond to the time intervals marked in red in the top panels.

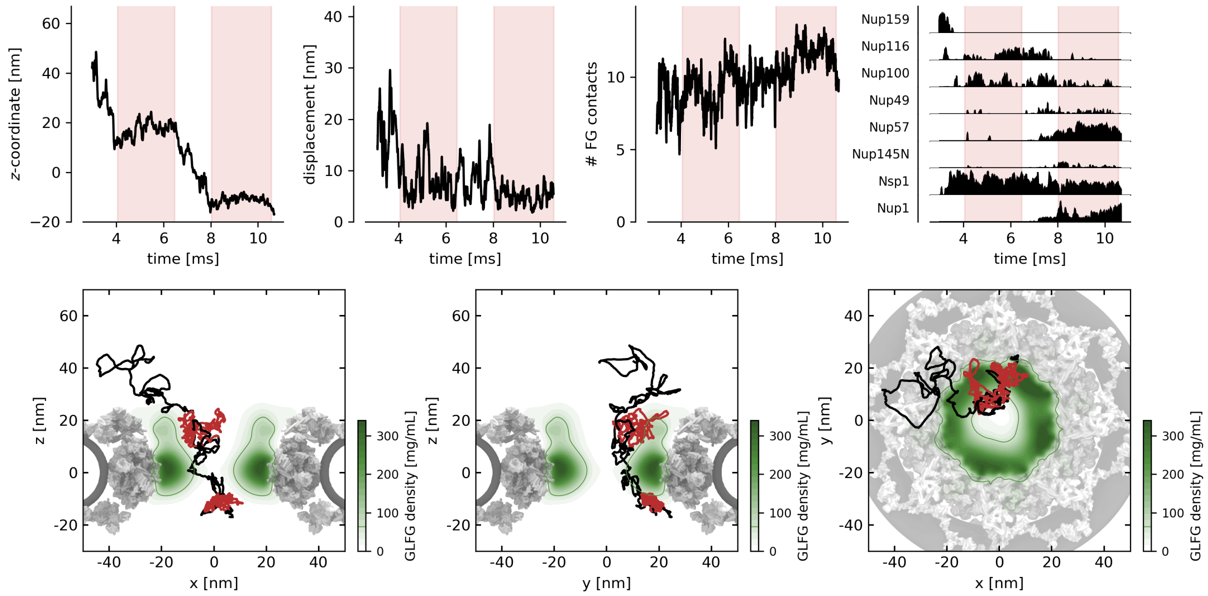

**Fig. S23. Detailed analysis of a single Kap95 translocation event (KAP #80)**. Top panels (left to right): (1) Axial position ($z$-coordinate) of the Kap95 as a function of time, showing its progression through the pore. (2) Instantaneous speed of the Kap, calculated as the displacement over 50 µs time windows. (3) Number of concurrent FG-motif interactions over time. (4) Breakdown of FG-contacts per FG-Nup type. In all four top panels, red intervals indicate dwell times within the FG-meshwork. Bottom panels (left to right): (1) 2D $x$–$z$ projection of the Kap trajectory, overlaid on the average density map of GLFG-Nups. (2) Same trajectory shown in the $y$–$z$ plane. (3) Same trajectory shown in the $x$–$y$ plane (top view of the pore). The red segments in the trajectory correspond to the time intervals marked in red in the top panels.

#### Supplementary Movies

**Movie S1. NPC with 80 Kaps.** A short trajectory of the coarse-grained 1BPA molecular dynamics simulation of the NPC with 80 Kaps.

**Movie S2. Trajectory of a mobile Kap translocating through the NPC.** A selected Kap overlayed on top of the FG-motif distribution (left) and the overall protein distribution of both FG-Nups and Kaps (right). The movie corresponds to the Kap shown in Fig. 4 of the main text.

**Movie S3. Trajectory of an immobile Kap in the NPC.** A selected Kap overlayed on top of the FG-motif distribution (left) and the overall protein distribution of both FG-Nups and Kaps (right).
